# The virulence regulator VirB from *Shigella flexneri* uses a CTP-dependent switch mechanism to activate gene expression

**DOI:** 10.1101/2023.06.01.543266

**Authors:** Sara Jakob, Wieland Steinchen, Juri Hanßmann, Julia Rosum, Manuel Osorio-Valeriano, Pietro I. Giammarinaro, Gert Bange, Martin Thanbichler

## Abstract

The transcriptional antisilencer VirB acts as a master regulator of virulence gene expression in the human pathogen *Shigella flexneri*. It binds defined sequences (*virS*) upstream of VirB-dependent promoters and counteracts their silencing by the nucleoid-organizing protein H-NS. However, its precise mode of action remains unclear. Notably, VirB is not a classical transcription factor but related to DNA partitioning proteins of the ParB family, which have recently been recognized as DNA-sliding clamps using CTP binding and hydrolysis to control their DNA entry gate. Here, we show that VirB binds CTP, embraces DNA in a clamp-like fashion upon its CTP-dependent loading at *virS* sites and slides laterally on DNA after clamp closure. Mutations that prevent CTP binding block the loading of VirB clamps in *vitro* and the formation of VirB nucleoprotein complexes *in vivo*. Thus, VirB represents a CTP-dependent molecular switch that uses a loading-and-sliding mechanism to control transcription during bacterial pathogenesis.

## Introduction

*Shigella* species are the causative agents of bacillary dysentery^1^ and the second leading cause of diarrheal mortality, with more than 200.000 deaths per year worldwide^2^. After ingestion, they invade the colonial epithelium, replicate in the cytoplasm of epithelial cells then propagate the infection by spreading from cell to cell, driven by actin-based motility. These processes are facilitated by a multitude of virulence proteins, known as effectors, which are produced in two consecutive waves and directly injected into the host cytoplasm by means of a type III secretion system (T3SS)^3^. The T3SS apparatus as well as most effectors are encoded on a large (∼220 kb) virulence plasmid, named pINV^4–6^. Many of the pINV-associated genes are controlled at the transcriptional levels by a three-tiered regulatory cascade^7^ that is triggered upon transition of *Shigella* cells to human body temperature (37 °C)^8^. It initiates with the production of the transcriptional activator VirF^9^, driven by temperature-induced changes in the structure of the *virF* promoter region^10–14^. Among the regulatory targets of VirF is the gene for the second-tier regulator VirB^15–17^, which activates the expression of ∼50 genes, coding for the structural components of the T3SS and for effectors mediating host invasion^18^. The VirB regulon also includes the gene for the third-tier regulator MxiE^18^, which later promotes the production of effectors involved in post-invasion processes, including the subversion of host immune response, innate immunity and cell death pathways^1, 7^.

Due to its central role in virulence gene expression, VirB is critical for pathogenicity in *Shigella* and mutants lacking this regulator are avirulent^17, 19, 20^. Interestingly, while VirF and MxiE are classical AraC-type transcriptional activators^9, 21, 22^, VirB is an atypical transcription factor related to the ParB family of DNA partitioning proteins^23–26^. Members of this family usually function in the context of ParAB*S* DNA segregation systems, where they serve as centromere-binding proteins that interact dynamically with the partition ATPase ParA to mediate the separation of newly replicated sister chromosomes or low-copy number plasmids^27^. ParB proteins form dimers and interact specifically with short (16 bp) palindromic DNA sequences called *parS* that are clustered close to the replication origin of target molecules^28, 29^. After initial specific binding to individual *parS* sites, they spread laterally into the neighboring DNA segments, forming a large nucleoprotein assembly (partition complex) that typically includes 10-20 kb of the origin region^30–33^. Recent work has clarified the mechanistic basis of the spreading process by showing that ParB proteins constitute a novel class of molecular switches and act as DNA-sliding clamps that use cytidine triphosphate (CTP) binding and hydrolysis to control the opening state of their DNA entry gate ^34–39^. The CTP-binding site is located in the N-terminal ParB/Sulfiredoxin (Srx) domain of ParB^38, 39^, which is followed by a central helix-turn-helix (HTH) *parS*-binding domain^40–43^, a non-structured linker region and a C-terminal dimerization domain, which tightly links the two subunits of a ParB dimer^41, 44^. Upon juxtaposition of the HTH domains on a palindromic *parS* site, the two N-terminal ParB/Srx domains associate in a CTP-dependent manner and thus close the ParB dimer into a ring-like structure, with bound DNA entrapped in between the two subunits^35, 37, 39^. Ring closure releases the HTH domains from *parS* and thus enables the ParB clamp to slide into the flanking DNA regions, making *parS* accessible again to other ParB dimers^35, 39, 45, 46^. The slow intrinsic CTPase activity of ParB rings eventually leads to CTP hydrolysis, triggering clamp opening and the dissociation of ParB from the DNA^34, 36, 37^. Spontaneous nucleotide exchange then initiates the next loading cycle. This sequence of events leads to the establishment of a 1D diffusion gradient of ParB clamps within the origin region, originating at *parS* clusters and shaped by the CTPase activity of ParB.

Intriguingly, the regulatory activity of VirB depends on its interaction with *parS*-like palindromic sequences (henceforth called *virS*) in the target promoter regions^23, 25, 26, 47–49^. These recognition sites can be located more than 1 kb upstream of the corresponding transcriptional start sites^48, 50^, indicating that VirB does not stimulate gene expression through a direct interaction with RNA polymerase. A series of studies showed that VirB indeed acts at a distance and counteracts the silencing of target promoters by the histone-like nucleoid-structuring protein H-NS, which associates with AT-rich regions in the surroundings or down-stream of *virS* sites^26, 48–52^. This anti-silencing mechanism was proposed to involve the spreading of VirB from its DNA recognition site, because the insertion of a roadblock between *virS* and the H-NS binding regions abolished the regulatory effect of VirB^49^. Consistent with this notion, recent work has shown that the *virS*-dependent interaction of VirB with promoter regions causes large-scale changes in DNA topology, leading to a local decrease in negative supercoiling that alleviates H-NS-mediated gene silencing^53^. However, the underlying mechanism remains unclear.

The homology of VirB to ParB proteins, its dependence on a *parS*-like binding site and its potential spreading within promoter regions raise the possibility that VirB could use a clamping-and-sliding mechanism similar to that reported for ParB to associate with its target DNA. In this study, we use a combination of structural modeling, molecular interaction studies, hydrogen-deuterium exchange mass spectrometry and *in vivo* localization studies to dissect the function of VirB. We demonstrate that VirB indeed binds CTP but appears to lack appreciable CTPase activity. Consistent with previous results^25, 54^, we show that VirB has strong non-specific DNA-binding activity in standard buffers. However, under more stringent conditions, it accumulates on DNA in a strictly *virS* and CTP-dependent manner, with *virS* acting as an entry site that mediates the loading of multiple VirB dimers, which diffuse away from *virS* after the loading step. We further observe that the two N-terminal nucleotide-binding domains of VirB homodimerize in the presence of CTP, in a process stimulated by *virS* DNA. Finally, we show that CTP binding is essential for the formation of VirB nucleoprotein complexes *in vitro.* Together, these results indicate that VirB constitutes a distinct class of CTP-dependent molecular switches that use a loading-and-sliding mechanism control gene expression during bacterial pathogenesis.

## Results

### VirB is a homolog of plasmid-encoded ParB proteins with CTP-binding activity

Previous work has revealed a significant similarity of VirB to plasmid-encoded ParB proteins in terms of its amino acid sequence and domain architecture as well as its ability to interact specifically with a *parS*-like sequence motif^23, 24, 26^. Moreover, crystallographic studies showed that the *virS*-binding domain of VirB is structurally related to the HTH-domains of the ParB homologs ParB, SopB and KorB, encoded by the *E. coli* plasmids P1, F and RP4, respectively^25^. Prompted by these findings, we aimed to investigate whether VirB also shared the ability of ParB proteins to bind and hydrolyze CTP. To this end, we first generated an amino acid sequence alignment comparing the N-terminal domain of VirB with the corresponding N-terminal nucleotide-binding domain of various well-characterized ParB homologs from plasmid or chromosome partitioning systems. Importantly, three regions that critically contribute to CTP binding in ParB are also conserved in VirB (**Figure 1A**), including the C-motif, the GxRR-motif and the P-loop, which coordinate the cytidine base, the triphosphate moiety and the γ-phosphate group of the CTP nucleotide, respectively^36, 37, 39^. However, VirB features only one out of the two conserved residues required for CTP hydrolysis in chromosomally encoded ParB proteins^34, 36, 37^ (**Figure 1A**), suggesting that it may be able to bind CTP but potentially lack CTPase activity.

**Figure 1.**
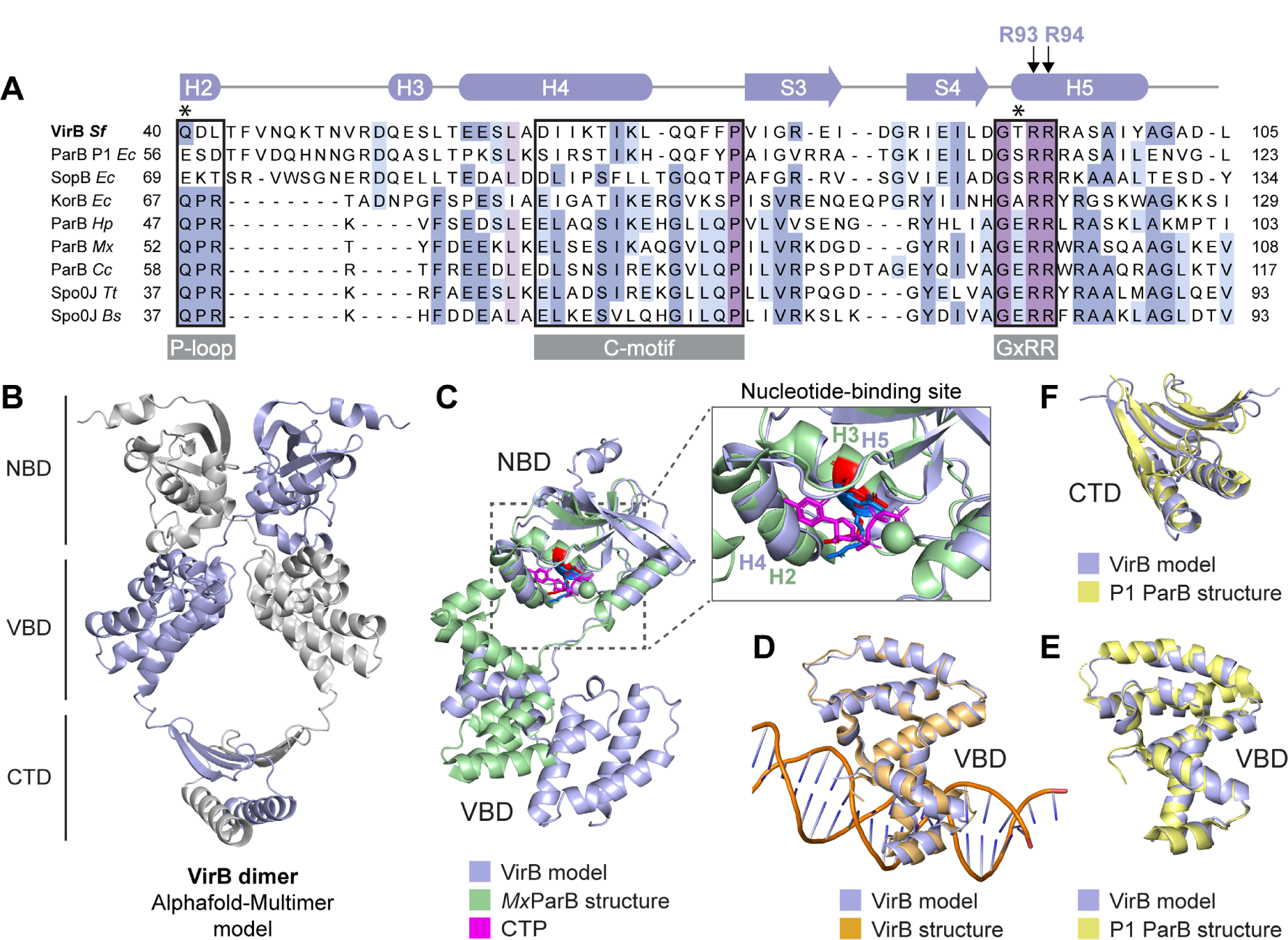
VirB is structurally related to plasmid-encoded ParB proteins. **(A)** Multiple sequence alignment comparing VirB from *S. flexneri* with plasmid- and chromosomally encoded ParB orthologs, with a focus on the nucleotide-binding region. Conserved motifs with critical roles in nucleotide binding, including the P-loop, the C-motif and GxRR motif^37^, are highlighted by black frames. The schematic at the top represents the predicted secondary structure of VirB. Two conserved arginine residues (R93 and R94) in the GxRR motif, which have been shown to be essential for nucleotide binding in other family members, are marked. Asterisk indicate the residues that are were shown to be critical for CTP hydrolysis in chromosomally encoded ParB proteins^34^, ^36, 37^. The proteins aligned and their UniProt accession numbers are VirB of *S. flexneri* (P0A247), ParB from *Escherichia coli* plasmid P1 (P07621), SopB from *E. coli* plasmid F (P62558); KorB from *E. coli* plasmid RK2 (P07674), ParB from *Helicobacter pylori* (O25758), ParB from *Myxococcus xanthus* (Q1CVJ4), ParB from *Caulobacter crescentus* (P0CAV8), Spo0J from *Thermos thermophilus* (Q72H91) and Spo0J from *Bacillus subtilis* (P26497). **(B)** Structural model of an *S. flexneri* VirB dimer, generated with AlphaFold-Multimer^55^. The nucleotide-binding domain (NBD), the *virS*-binding domain (VBD) and the C-terminal dimerization domain (CTD) are indicated. **(C)** Overlay of the predicted structure of the NBD of VirB with the crystal structure of the NBD of *M. xanthus* ParB (*Mx*ParB) (PDB: 7BNR), with an RMSD of 1.512 Å for 45 aligned C_α_ atoms. Close-up of the nucleotide-binding site, with the conserved arginine residues in the GxRR motif of VirB (R93 and R94 in helix H5) shown in blue and their equivalent residues (R94 and R95 in helix H3) in *Mx*ParB shown in red. **(D)** Overlay of the modeled VBD of VirB with the crystal structure of the VBD bound to the *virS* site of the *S. flexneri icsB* gene (PDB: 3W3C), with an RMSD of 0.883 Å for 98 aligned C_α_ atoms. **(E)** Overlay of the modeled VBD of VirB with the crystal structure of the DNA-bound HTH-domain of *E. coli* plasmid P1 ParB (PDB: 1ZX4), with an RMSD of 0.811 Å for 93 aligned C_α_ atoms. **(F)** Overlay of the CTD of modeled VirB with the corresponding domain of *E. coli* plasmid P1 ParB (PDB: 1ZX4), with an RMSD of 1.274 Å for 90 aligned C_α_ atoms.

To further analyze the function of VirB, we generated a structural model of a VirB homodimer, using AlphaFold-Multimer^55^ (**Figure 1B**). This model suggests that VirB can form a clamp-like structure similar to that reported for ParB^36, 37, 39^, with the two subunits interacting in the N-terminal and C-terminal regions. The center of the closed dimer features an opening flanked by non-structured linker regions that is large enough to accommodate a DNA molecule and lined by an abundance of positively charged residues (**Figure S1**). As in ParB, the interaction in the N-terminal regions is predicted to rely on the homodimerization of the two ParB/Srx nucleotide-binding domains (NBDs), which involves a crossover of the two polypeptide chains that places the NBD of each *cis*-subunit next to the *virS*-binding domain (VBD) of the *trans*-subunit. A superimposition of the structural model of VirB with the crystal structure of the chromosomally encoded ParB homolog from *Myxococcus xanthus*^37^ indicates that the NBDs of the two proteins have a similar fold, especially in regions forming the nucleotide-binding pocket of ParB, with a root-mean-square deviation (RMSD) of 1.51 Å for 45 paired C_α_ atoms (**Figure 1C**). However, VirB is distinguished from ParB by an N-terminal helix as well as two adjacent long antiparallel β-strands that appear to connect the nucle-otide-binding region of VirB to an opposing α-helix (corresponding to helix 4 of ParB), thereby potentially reducing the conformational flexibility of the NBD. The predicted structure of the VBD is highly similar to a previously solved crystal structure of the VBD in complex with *virS* DNA ^25^ (RMSD of 0.883 Å for 98 paired C_α_ atoms) (**Figure 1D**), which underscores the validity the modeling approach (**Figure 1D**). Moreover, while the VBD is structurally different from the HTH-domains of chromosomally encoded ParB homologs (**Figure 1C**), it shows striking similarity to the HTH-domain of ParB from *E. coli* plasmid P1 (**Figure 1D**). Considerable structural similarity is also observed for the C-terminal dimerization domains (CTDs) of VirB and P1 ParB (**Figure 1E**), supporting the notion that VirB has evolved from plasmid partitioning proteins.

### VirB has CTP-binding activity but lacks CTPase activity

Since key features of the nucleotide-binding pocket of ParB were conserved in VirB, we aimed to investigate whether VirB was also able to interact with CTP, using quantitative nucleotide binding assays based on isothermal titration calorimetry (ITC) (**Figure 2A**). First, we analyzed the binding of the poorly hydrolyzable CTP analog CTPγS, which was chosen to avoid potential adverse effects of CTP hydrolysis on the measurements. We observed that VirB bound this nucleotide with an affinity (*K*_D_ ∼ 12 µM) that is comparable to the one measured for ParB^34, 37^ and high enough to ensure its saturation under *in vivo* conditions^56^. We also observed an interaction of VirB with cytidine diphosphate (CDP), although the binding affinity for this nucleotide was more than 7-fold lower (*K*_D_ ∼ 91 µM). Similar results were obtained using microscale thermophoresis as an alternative technique to study the nucleotide-binding behavior of VirB (**Figure 2B** and **Figure S2**). Notably, CTP binding was drastically reduced upon the mutation of arginine residues (R93, R94) in the conserved GxRR motif to alanines (see **Figure 1A**), suggesting that VirB and ParB share a similar mode of nucleotide binding. Given that some proteins, such as sulfiredoxins^57, 58^ and free-serine kinases^59, 60^, contain ParB/Srx-like domains with TP-binding activity, we also tested VirB for its ability to interact with ATP. However, no significant binding was observed (**Figure S2**).

**Figure 2.**
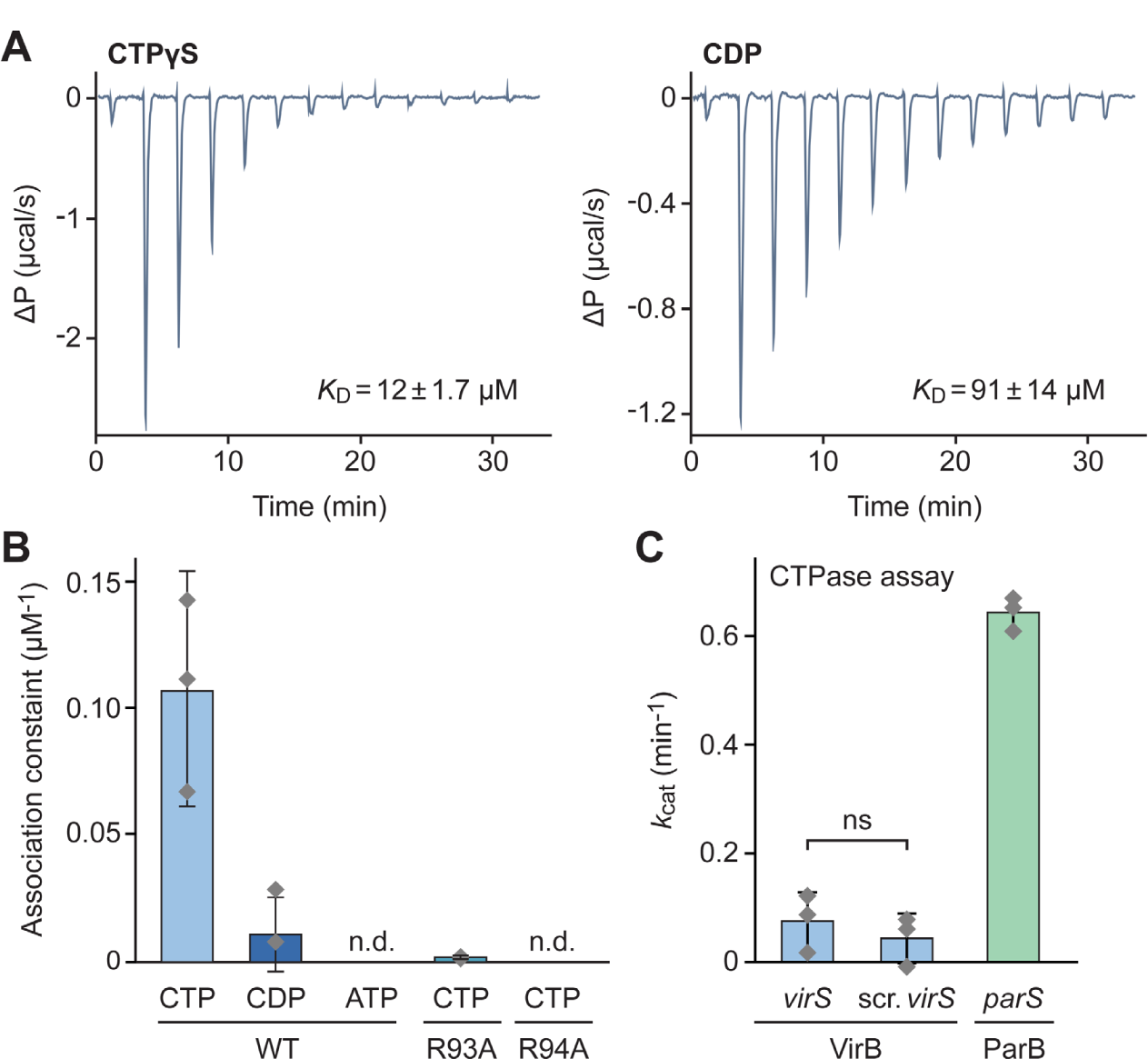
VirB is a CTP-binding protein. **(A)** Isothermal titration calorimetry analysis of the interaction of VirB with CTPγS and CDP. A solution of VirB (115 µM) was titrated with a stock solution (1.55 mM) of the indicated nucleotides. The graphs indicate the heat changes observed after each of the 13 injections. The *K*_D_ values obtained are given in the graphs. **(B)** Microscale thermophoresis analysis of the interaction of VirB and its R93A and R94A variants with CTP, CDP and ATP. The diamonds show the mean equilibrium association constants (1/*K*_D_) obtained for the indicated conditions (± SD; n=2-3 independent experiments, each performed in triplicate). The bars give the values determined by fitting the combined results of all nine replicates. The underlying titration curves are given in **Figure S2**. The corresponding *K*_D_ values are: WT-CTP (9.3 µM), WT-CDP (99 µM) and R93A-CTP (981 µM). n.d. = not detectable. **(C)** CTPase activities of VirB and *Mx*ParB. VirB or *Mx*ParB (5 µM) were incubated with 1 mM CTP in the presence of double-stranded DNA oligonucleotides containing a *virS* (0.5 µM), a scrambled *virS* (0.5 µM) or an *M. xanthus parS* (0.3 µM) motif. CTP hydrolysis rates were determined using an NADH-coupled enzyme assay. The columns indicate the mean (± SD) of three independent measurements (represented by diamonds).

We then went on to clarify whether VirB was able to hydrolyze the bound CTP. In the case of ParB, significant CTPase activity is only detectable when the protein is incubated with both nucleotide and *parS* DNA. However, VirB did not hydrolyze CTP under any of the condition tested, even if *virS* DNA was included in the reaction (**Figure 2C**). These results indicate that VirB binds CTP but lacks appreciable CTPase activity. However, it is possible that additional, thus-far unknown factors are required to trigger the hydrolytic reaction.

### VirB is loaded onto DNA in a CTP- and *virS*-dependent manner

VirB was shown to specifically interact with *virS in vitro* and to require the presence of *virS* upstream of target promoters for its regulatory activity *in vivo*^25, 26, 48, 49, 51, 53^. To clarify the role of CTP-binding in the interaction of VirB with target DNA regions, we analyzed its DNA-binding behavior using a previously established biolayer interferometry assay^35^. For this purpose, DNA fragments (215 bp) that contained the *virS* sequence located upstream of the *S. flexneri icsB* promoter^26, 47^ as well as its flanking regions were immobilized on biosensors such that both of their ends were stably linked to the sensor surface (**Figure 3A**). Subsequently, these closed fragments were probed with VirB in the absence or presence of CTP. In both conditions, VirB showed strong DNA-binding activity. However, nucleotide-free reactions reproducibly yielded biphasic association curves when VirB was used at elevated concentrations, with the signal decreasing markedly during the loading phase (**Figure 3B**). In the presence of CTP, by contrast, VirB stably associated with the DNA in all cases (**Figure 3C**). However, almost identical results were obtained with closed DNA fragments lacking a *virS* sequence (**Figure S3**). Similarly, VirB also showed a strong *virS*-independent interaction with biosensors carrying open double-stranded oligonucleotides, although the presence of CTP again appeared to modulate its binding behavior (**Figure S4**). These findings imply that VirB has strong non-specific DNA-binding activity, consistent with the high density of positive charges in its VBD and CTD (**Figure S1**). This property was likely to obscure the specific effects that *virS* and CTP might have on the DNA-binding behavior of VirB under the conditions used (150 mM NaCl) *in vitro*.

**Figure 3.**
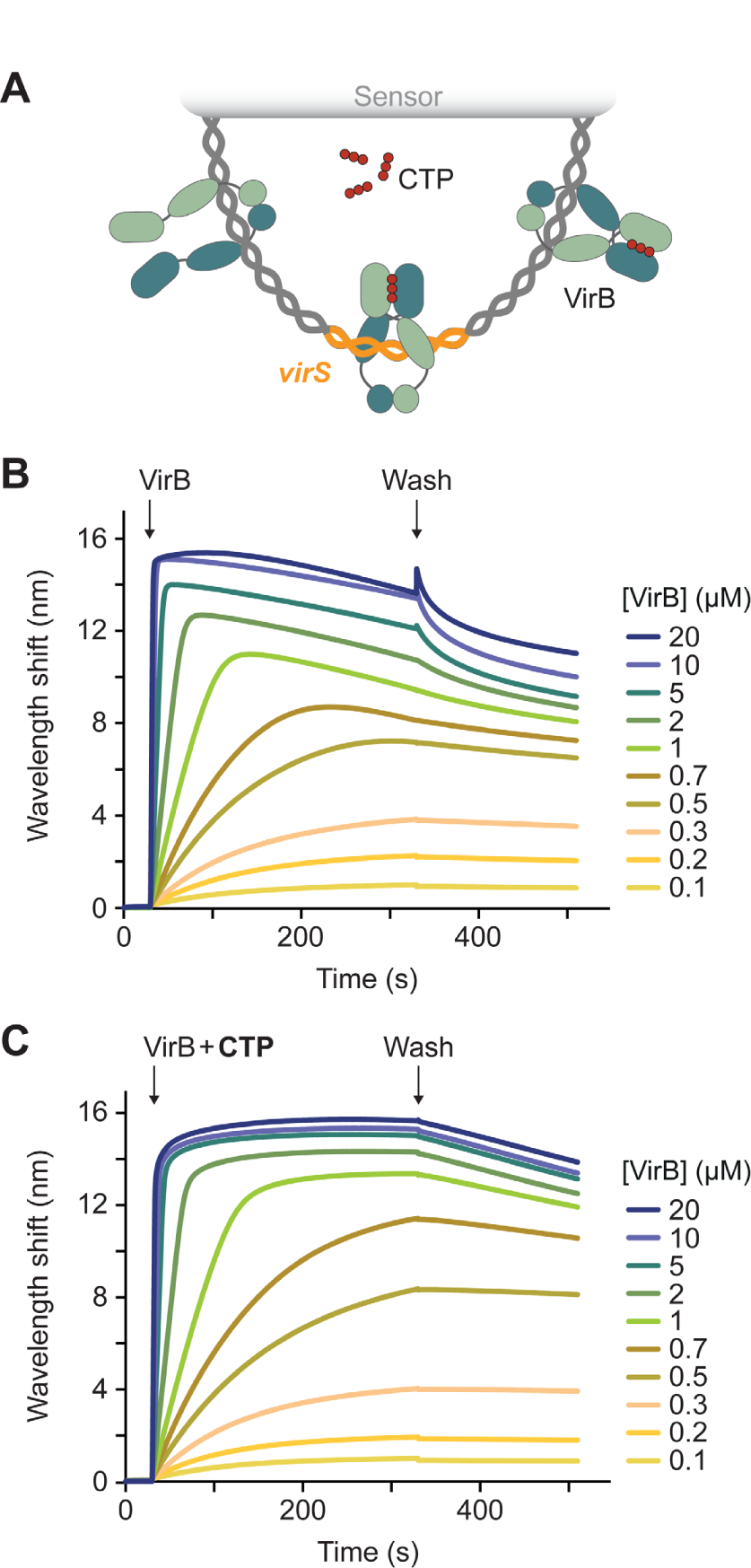
CTP-binding modulates the DNA-binding activity of VirB in low-stringency conditions. **(A)** Schematic of the biolayer interferometry (BLI) setup used for the analyses in panels B and C. A double-biotinylated dsDNA fragment (215 bp) containing a central *virS* sequence (in orange) was immobilized on a streptavidin-coated biosensor and probed with VirB (green). **(B,C)** BLI analysis of the DNA-binding behavior of VirB in the (B) absence and (C) presence of CTP (1 mM) in low-stringency buffer (150 mM NaCl). Biosensors carrying the target DNA (at a density corresponding to a wavelength shift of ∼1.3 nm) were probed with the indicated concentrations of VirB. At the end of the association reactions, the biosensors were transferred into protein- and nucleotide-free buffer to monitor the dissociation reactions (wash). The graphs show the results of a representative experiment (n=3 independent replicates).

It was possible that more stringent conditions were required to discriminate between specific and non-specific interactions of VirB with its target DNA. We therefore repeated the binding assays at elevated salt concentrations (500 mM NaCl) to weaken electrostatic interactions between positively charged residues and the DNA phosphate backbone. Importantly, in the modified buffer, VirB no longer displayed any non-specific DNA-binding activity and only showed a marginal association with closed, *virS*-containing DNA when assayed in the absence of CTP. By contrast, strong binding was observed if CTP was included in the reaction (**Figure 4A**). In line with the results of the nucleotide binding assays (**Figure 2**), this interaction was strictly dependent on CTP and not observed in assays using CDP, UTP, ATP or GTP instead (**Figure S5**). Moreover, it required the presence of *virS*, because only residual binding was detected for similar DNA fragments lacking a *virS* sequence (**Figure 4B**). A titration analysis showed that the CTP-dependent binding of VirB to *virS*-containing DNA occurred with high affinity (apparent *K*_D_ = 2.3 µM), with more than 14 dimers accumulating on each DNA molecule at saturation (**Figure 4C**). This behavior is highly reminiscent of the CTP-dependent loading of ParB onto *parS*-containing centromeric DNA, suggesting that VirB could use a similar loading-and-sliding mechanism to accumulate in promoter regions.

**Figure 4.**
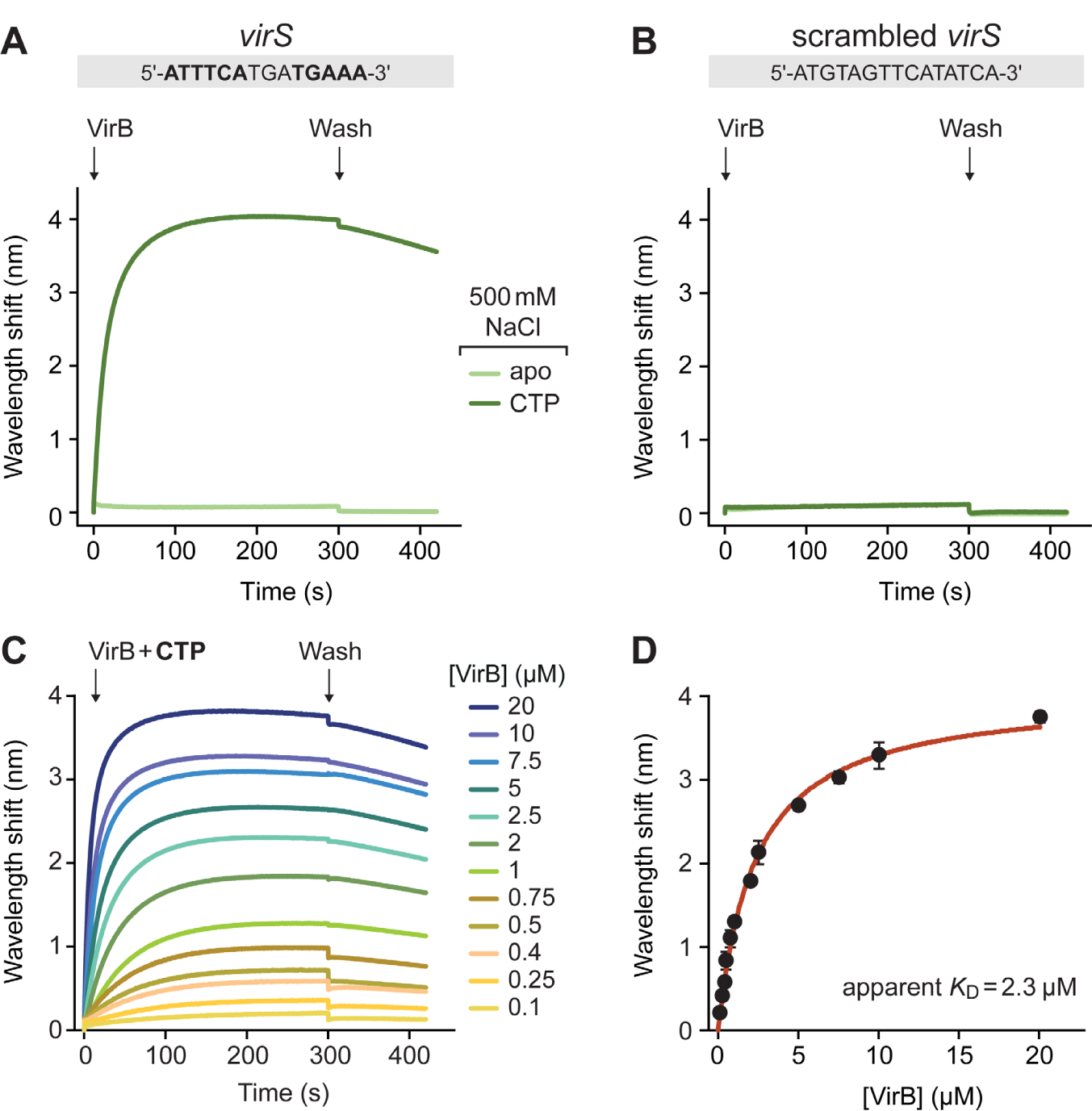
VirB requires CTP- and *virS* binding to accumulate on DNA in high-stringency conditions. **(A)** Biolayer interferometry analysis of the interaction of VirB with *virS*-containing DNA (215 bp) in high-stringency buffer (500 mM NaCl). Biosensors carrying a double-biotinylated, *virS*-containing DNA fragment (at a density corresponding to a wavelength shift of ∼0.5) were probed with VirB (20 µM) in the absence or presence of CTP (1 mM). The *virS* sequence used is shown above the graph. **(B)** Same as in panel A, using a DNA fragment with a scrambled *virS* site. **(C)** Titration of double-biotinylated *virS*-containing DNA (215 bp) with increasing concentrations of VirB in the presence of CTP (1 mM) in high-stringency buffer (500 mM NaCl). DNA was immobilized as described in panel A. **(D)** DNA-binding affinity of VirB in high-stringency conditions. The maximal wavelength shifts measured at equilibrium in the traces shown in panel C were plotted against the corresponding VirB concentrations. Error bars indicate the SD (n=3 independent replicates). A one-site specific-binding model was used to fit the data. The calculated *K*_D_ value is given in the graph. Note that the wavelength shift observed is directly proportional to the amount of matter associated with the biosensor surface. Based on the molecular weights of the immobilized DNA fragment (134 kDa) and the VirB dimer (71 kDa), the signals obtained indicate the binding of ∼14 VirB dimers at saturating concentrations.

### VirB forms DNA-sliding clamps

Previous work has shown that ParB clamps cannot stably associate with *parS*-containing DNA molecules that have open ends, because they slide off the DNA as soon as they are released from their loading site^35, 39^. To clarify the state of VirB after its CTP-dependent loading at *virS*, we therefore performed bio-layer interferometry analyses of its interaction with open DNA fragments, again using stringent conditions that prevented non-specific DNA binding. For this purpose, we first probed biosensors carrying short (23 bp) double-stranded oligonucleotides with a central *virS* site that were only attached at one of their ends, so that the other end remained open (**Figure 5A**). Subsequently, we analyzed for the interaction of VirB with the immobilized DNA in the presence of CTP. Notably, even at very high concentrations, VirB barely associated with the open target DNA (**Figure 5B**), even though the same oligonucleotides were densely covered with VirB in low-stringency conditions, which verifies the functionality of the biosensors used (**Figure S4**). To further test for the ability of VirB to slide on DNA, we compared the interaction of VirB with biosensors carrying closed *virS*-containing DNA fragments (215 bp) before and after cleavage of the fragments with the restriction endonuclease NdeI (**Figure 5C**). As expected, VirB strongly accumulated on closed target molecules when provided with CTP. However, after opening of the fragments by NdeI treatment, the maximum binding levels were approximately fourfold lower, suggesting that VirB dimers are lost from the DNA after binding if its ends are no longer blocked (**Figure 5D**). Moreover, during the association phase, the signal increased only briefly and then started to decrease, almost returning back to the baseline level. This behavior may be caused by the continuous loading and closure of VirB clamps at *virS*, which slide off the DNA and remain closed thereafter, leading to a steady decrease in the concentration of binding-competent, open VirB dimers.

**Figure 5.**
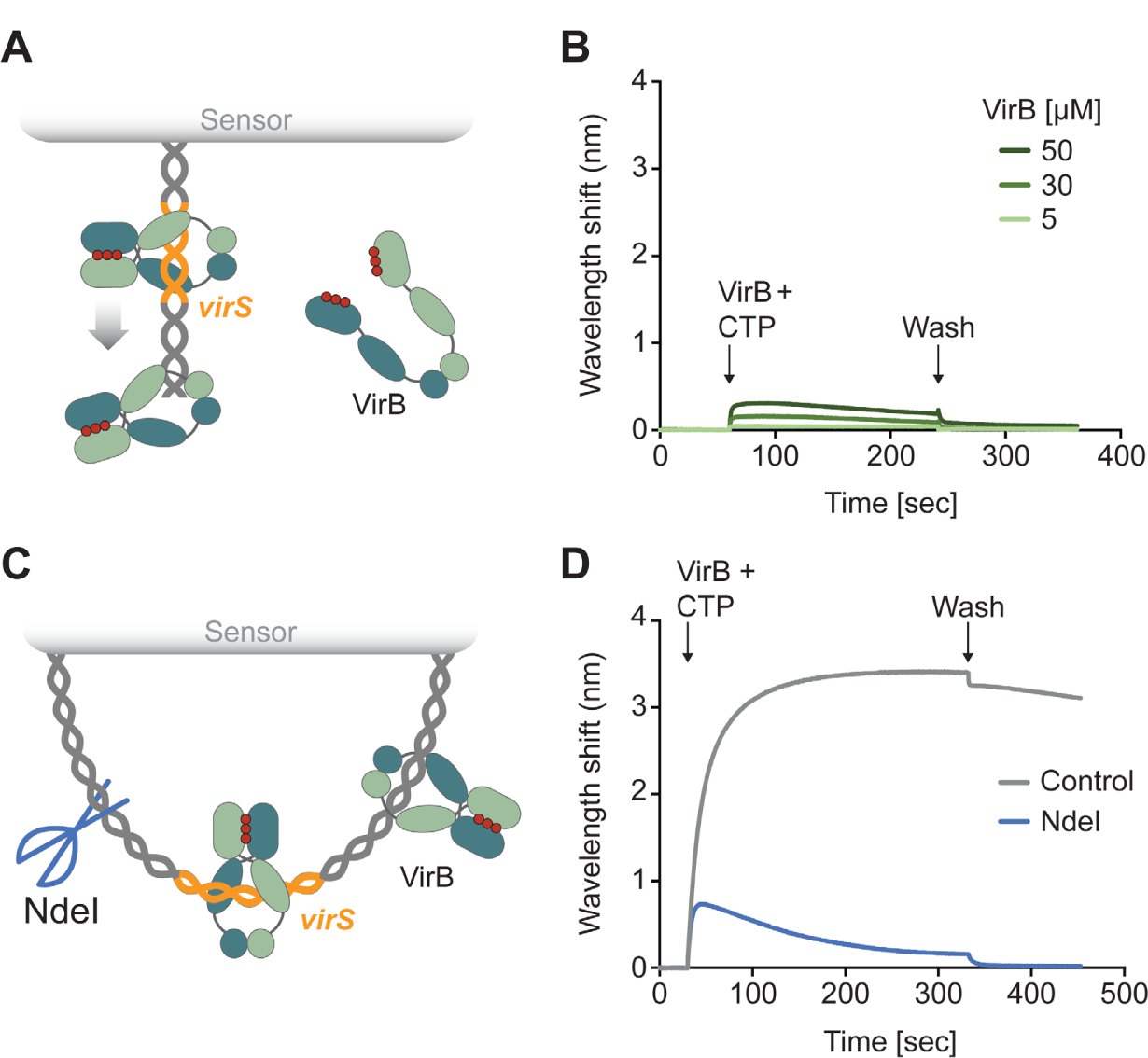
VirB requires CTP- and *virS* binding to accumulate on DNA in high-stringency conditions. **(A)** Biolayer interferometry (BLI) setup used to analyze the interaction of VirB with short, open *virS* DNA. A double-stranded *virS*-containing oligonucleotide biotinylated at one of its ends was immobilized on a streptavidin-coated biosensor. **(B)** BLI analysis using the setup described in panel A. The biosensors were probed with VirB at the indicated concentrations in the presence of CTP (1 mM), using high-stringency buffer (500 mM NaCl). **(C)** BLI setup used to compare the interaction of VirB with closed and open *virS* DNA. A double-biotinylated *virS*-containing DNA fragment (215 bp) was immobilized on two streptavidin-coated biosensors. Prior to the BLI assay, the biosensors either treated with the restriction endonuclease NdeI to open the immobilized DNA or incubated in the absence of *Nde*I as a control. **(D)** BLI analysis using the setup described in panel C. The biosensors incubated with or without NdeI were probed with VirB (20 µM) in the presence of CTP (1 mM), using high-stringency buffer (500 mM NaCl). The graphs in panels B and D show the results of representative experiments (n=3 independent replicates each).

The DNA-binding behavior of VirB suggested that it used a loading-and-sliding mechanism analogous to the one reported for ParB. We therefore aimed to determine whether the CTP-dependent interaction of VirB with *virS*-containing DNA could trigger the homodimerization of its two NBDs and thus close the VirB dimer into a ring-like structure, as also suggested by the structural model (**Figure 1B**). For this purpose, we generated a mutant variant of VirB (VirB-C5S/Q15C) that lacked the native cysteine residue at position 5 and carried an engineered cysteine residue at position 15, adjacent to the symmetry axis of the closed complex. (**Figure 6A**). Upon ring closure, the newly introduced C15 residues in the two NBDs were placed next to each other, enabling their covalent crosslinking by the bifunctional thiol-reactive compound bis-maleimidoethane (BMOE). Using this approach, we observed that most VirB dimers remained in the open state when incubated alone or in the sole presence of *virS* DNA. However, upon the addition of CTP and, even more so, a combination of CTP and *virS* DNA, the proportion of closed complexes increased considerably (**Figure 6B,C**). These findings support a model in which VirB forms DNA-sliding clamps that are loaded at *virS* sites and then closed by CTP-dependent homodimerization of the two NBDs.

**Figure 6.**
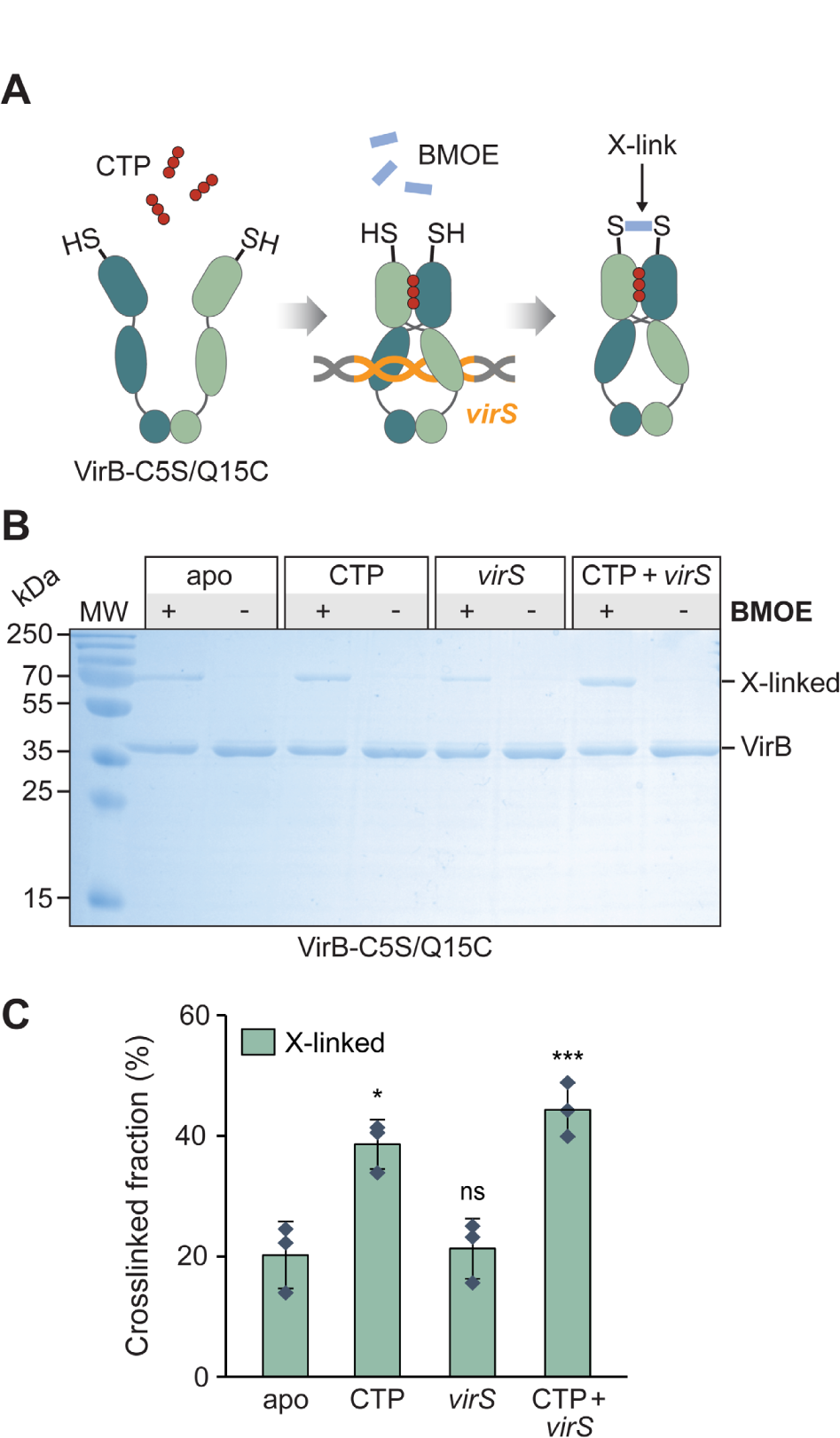
VirB clamps close in the presence of CTP and *virS*-containing DNA *in vitro*. **(A)** Schematic showing the crosslinking assay used to detect VirB clamp closure. The engineered C15 residue of VirB-C5S/Q15C is shown as a thiol group (-SH). The closure of the VirB clamp reduces the distance between the two C15 residues, thereby enabling their covalent crosslinking by the bifunctional thiol-reactive crosslinking agent BMOE. **(B)** SDS-gel showing the protein species obtained in the *in vitro* crosslinking analysis. VirB-C5S/Q15C was incubated for 5 min alone, with CTP (1 mM), with a double-stranded DNA oligonucleotide containing a *virS* motif (1 µM; virS-icsB-for/virS-icsB-rev) or with both CTP and *virS* DNA prior to crosslinking with BMOE and analysis of the reaction products by SDS-PAGE. Monomeric VirB and the dimeric crosslinking product (X-linked) are indicated. MW: Molecular weight marker. **(C)** Quantification of the fractions of crosslinked protein obtained in the indicated conditions. The columns display the mean (±SD) of three independent measurements (diamonds). *p<0.05, ***p<0.005, ns: not significant (Welch’s t-test; compared to the apo state).

Interestingly, similar results were obtained when the assay was performed with the wild-type protein, exploiting the native C5 residue for the crosslinking reaction (**Figure S6**). In this case, crosslinking was only mildly stimulated by CTP and largely dependent on the presence of *virS*. According to the structural model, C5 is located in N-terminal helix of VirB, which is predicted to closely associate with the remaining part of the NBD. This arrangement would place the two C5 residues in the VirB dimer at a distance (39 Å) too large to allow their crosslinking by BMOE (8 Å length) (**Figure S6A**). The high efficiency of the crosslinking reaction in the presence of CTP and *virS* could thus indicate that the N-terminal helices are released upon homodimerization of the NBDs.

### CTP and *virS* DNA regulate the dynamics of VirB clamp closure

To further investigate the role of CTP and *virS* binding in the closure of VirB clamps, we analyzed the structural dynamics of VirB using hydrogen-deuterium exchange (HDX) mass spectrometry, a technique that detects local changes in the accessibility of backbone amide hydrogens caused by conformational changes or ligand binding^61^. The initial set of experiments was performed in low-stringency buffer (150 mM NaCl) (**Figures S7A and S8A**). Under these conditions, the addition of a double-stranded oligonucleotide containing a scrambled, non-functional *virS* sequence (**Figure 4B**) led to a significant reduction in HDX in the C-terminal helices of the VBD as well as in the CTD compared to apo-VirB (**Figure 7A,C**), consistent with the idea that these regions are responsible for the strong non-specific DNA-binding activity of VirB (compare **Figure S1**). The same regions exhibited reduced HDX in the presence of a *virS*-containing oligonucleotide, but in this case the changes in the VBD were significantly more pronounced (**Figure 7A,C** and **Figure S9A**). Moreover, *virS* DNA additionally induced a strong reduction in HDX in the N-terminal half of the VBD, harboring the HTH-motif responsible for *virS* recognition^25^, as well as at the homodimerization interface of the NBD. Nucleotide-content analysis verified that the purified protein used for this analysis did not contain CTP or CDP (**Figure S10**). The juxtaposition of the two VBDs at the inverted repeats constituting the *virS* sequence thus appears to promote face-to-face interactions between the two NBDs independently of the presence of CTP. Notably, in the presence of *virS*, some peptides (e.g., residues 65-70 and 126-132) of the NBD showed a bimodal distribution of peptide ion intensities in their mass spectra, likely reflecting two populations with disparate HDX rates. This observation suggests that *virS*-bound VirB dimers dynamically switch between the open and closed state (**Figure S11**). In reactions containing only CTP, the differences in HDX observed in the NBD were even more pronounced than in the apo-state and extended throughout the homodimerization interface and the nucleotide-binding pocket (**Figure 7A,C**). Moreover, we observed reduced HDX in regions of the VBDs that are predicted to interact with each other in the closed complex, consistent with the idea that CTP binding promotes the closure of the VirB clamp and that this process is accompanied by rearrangements in VBDs that are likely to affect their DNA-binding behavior. However, again, peptides from the NBD showed a bimodal behavior, suggesting that the two NBDs are not stably associated with each other in the sole presence of CTP (**Figure S11**). Unlike in the case of CTP, the addition of CDP produced only minor shifts in the HDX pattern of the nucleotide-binding pocket and the VBD (**Figure S9B**), as expected from its inability to promote the loading of VirB clamps onto DNA *in vitro* (**Figure S5**). Finally, in reactions containing both *virS* DNA and CTP, strongly reduced HDX was observed throughout all domains of VirB, including all regions that were affected in reactions containing either of the two ligands (**Figure 7A,C**), with a considerably higher amplitude of reduction than with CTP or *virS* alone. Moreover, in this case, all peptides that showed bimodal behavior with CTP as the only ligand were consistently shifted to the slow-exchanging state. In line with the crosslinking data (**Figure 6** and **Figure S6**), this observation indicates that a larger fraction of VirB dimers transitions to the ligand-bound closed state if both ligands are present. However, very similar results were also obtained for reactions containing both CTP and an oligonucleotide with a scrambled *virS* sequence (**Figure 7C** and **Figure S9**), suggesting that, in low-stringency conditions, non-specific DNA-binding may potentially also be sufficient to stimulate VirB clamp closure.

**Figure 7.**
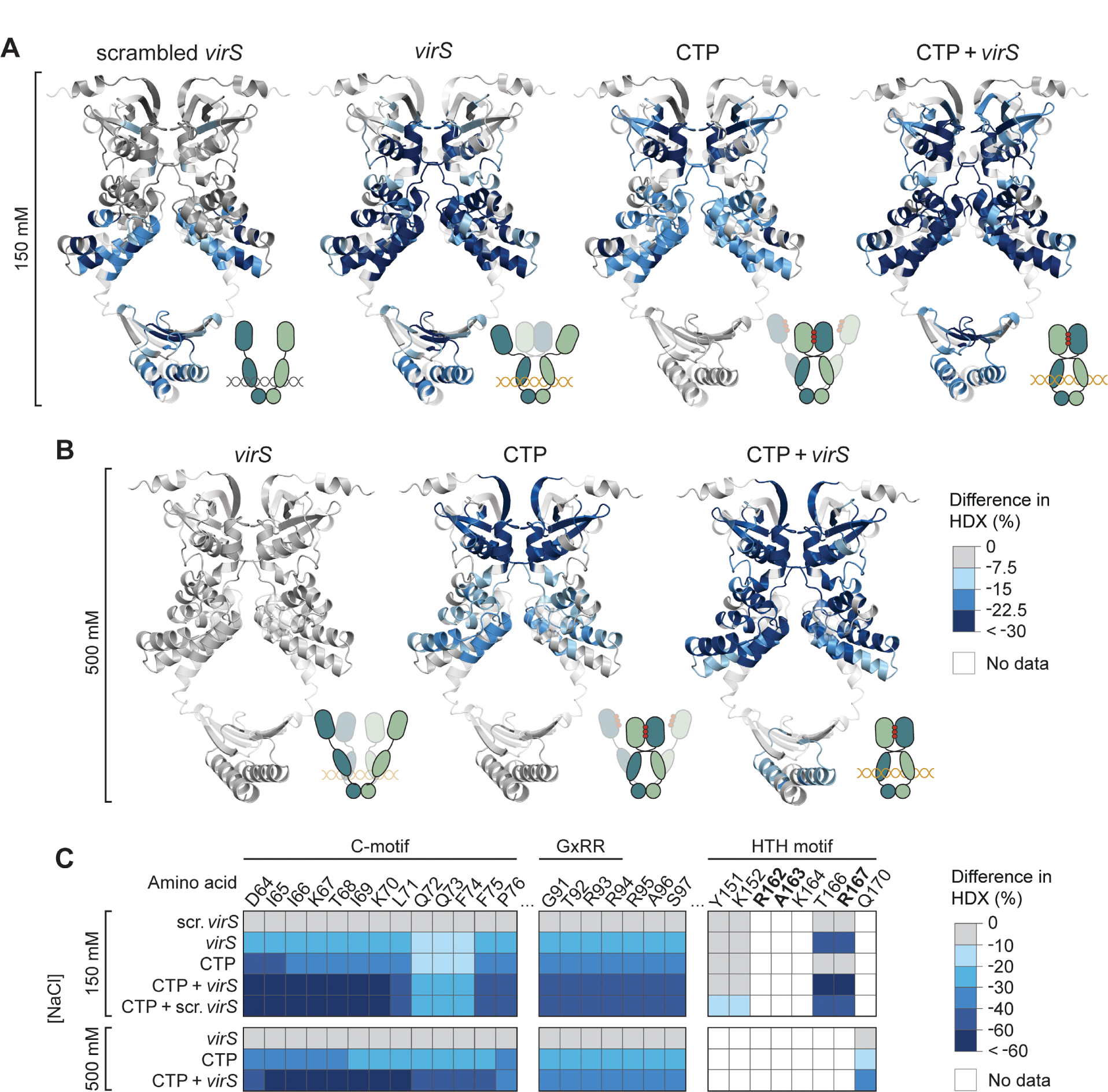
CTP and *virS* DNA cooperatively stimulate the homodimerization of the N-terminal region of VirB. **(A,B)** Hydrogen-deuterium exchange (HDX) mass spectrometry analysis of VirB in the presence of different ligands. VirB was incubated with an equimolar concentration of double-stranded DNA oligonuc-leotides containing a scrambled (scrambled-virS-for/scrambled-virS-rev) or intact (virS-icsB-for/virS-icsB-rev) *virS* motif and/or CTP (10 mM) in (A) low-stringency or (B) high-stringency buffer. Shown are the maximal differences in HDX obtained in the indicated conditions compared to the apo state, projected onto the AlphaFold-Multimer model of the VirB dimer. The color code is given in panel B. Blue color indicates regions that show reduced HDX upon ligand binding. The schematics next to the structural models indicate the most likely conformational state of the VirB dimer in the respective conditions. Protein regions not covered by any peptides are displayed in transparent white. **(C)** Heatmap of the maximal differences in HDX obtained in the indicated conditions for representative residues in the conserved C-, GxRR and HTH motifs of VirB. The color code is given on the right. A detailed report of the HDX analysis is given in **Data S1**.

Next, we performed the HDX analysis in high-stringency conditions (500 mM NaCl) that abolish non-specific DNA binding (**Figures S7B and S8B**). In this case, even *virS*-containing DNA did not produce a measurable change in the HDX pattern, although it mediated the robust loading of VirB onto closed DNA in the same buffer (see **Figure 4A**), indicating that *virS* binding was too dynamic to have a marked influence on the HDX reaction (**Figure 7B,C**). By, contrast, the addition of CTP again led to strong changes in the HDX pattern, similar to those observed in low-stringency conditions (**Figure 7B,C**). Again, several peptides in the NBD (residues 63-68, 79-75, and 118-128) and the VBD (residues 218-238) showed a bimodal behavior, suggesting that VirB alternated dynamically between the open and closed state in this condition (**Figure S12**). An additional, strong reduction in HDX was observed throughout all three domains of VirB when both CTP and *virS* were included in the reactions (**Figure 7B,C**), consistent with the finding that both ligands are required to trigger robust VirB clamp closure in high-stringency buffer (**Figures 4A,B and 6** and **Figure S6**).

### CTP binding is required for VirB to bind and regulate target promoters *in vivo*

Our analyses revealed that CTP was required to enable the specific loading of VirB clamps at *virS* sites in *vitro*. To clarify the role of CTP binding in the function of VirB *in vivo*, we made use of the R93A and R94A variants of VirB, which both lacked appreciable affinity for CTP (**Figure 2B**). Biolayer interferometry assays confirmed that these variants were no longer able to accumulate on closed *virS-*containing DNA in high-stringency conditions, indicative of a defect in CTP-mediated clamp closure (**Figure 8A**). Inspired by work in *S. flexneri*^62^, we then devised an *in vivo* assay that allowed us to visualize the recruitment of VirB to a *virS*-containing plasmid in the heterologous host *E. coli*, a species closely related to *S. flexneri*^63^ that has been commonly used to investigate the mechanistic basis of gene regulation by VirB^47, 49, 50^. To this end, *E. coli* was transformed with a low-copy plasmid carrying the upstream region (187 bp) of the *S. flexneri icsB* gene, including the previously reported *virS* site, which was shown to be essential for the VirB-dependent regulation of *icsB* expression *in vivo*^47^. The resulting strain was then additionally transformed with expression plasmids that allowed the production of fluorescently (mVenus-) tagged versions of VirB or its two mutant variants under the control of an arabinose-inducible promoter (**Figure 8B**). Upon induction, ∼80% of the cells producing the wild-type mVenus-VirB fusion formed one to several bright foci per cell (**Figure 8C,D**), reminiscent of results obtained previously for a synthetic *virS*-containing plasmid in pINV-free *S. flexneri* cells^62^. By contrast, only diffuse fluorescence was observed for cells producing the R93A or R94A variant, indicating that CTP binding is critical for the accumulation of VirB in the *icsB* promoter region *in vivo* (**Figure 8C,D**). Notably, the wild-type protein failed to form foci in cells that contained a low-copy plasmid lacking the *icsB* upstream region instead of the original construct, consistent with the notion that the CTP-dependent loading of VirB in target promoter regions is strictly dependent on the presence of *virS* (**Figure 8C,D**).

**Figure 8.**
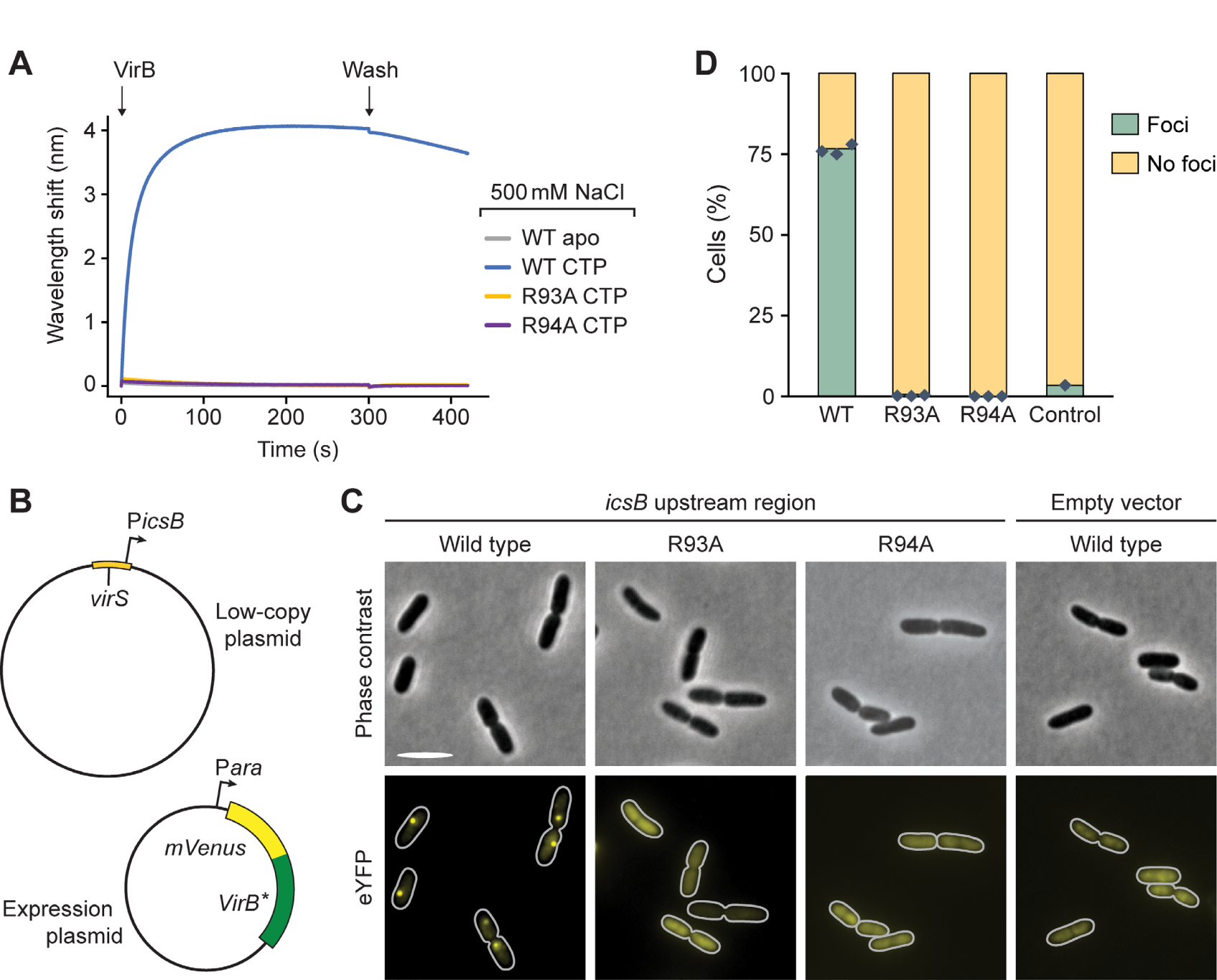
CTP-binding is critical for the loading of VirB on *virS*-containing DNA *in vivo*. **(A)** Biolayer interferometry analysis of the interaction of VirB-R93A and VirB-R94A with a closed, *virS*-containing DNA fragment (215 bp) in high-stringency buffer (500 mM NaCl). After derivatization with the double-biotinylated *virS* DNA, the biosensor was probed with wild-type VirB or its mutant derivatives (20 µM) in the absence (apo) or presence of CTP (1 mM). **(B)** Plasmids used for the *in vivo* binding assay. **(C)** Localization patterns of wild-type or mutant mVenus-VirB fusions in *E. coli* strains that harbor low-copy plasmids with or without the *icsB* upstream region. Cells carrying a low-copy containing (pSJ30) or lacking (pSJ31) the *icsB* upstream region were transformed with expression plasmids that allow the production of the indicated mVenus-VirB variants under the control of an arabinose-inducible promoter (pSJ18, pSJ20, pSJ21). Transformants were induced with 0.1% arabinose for 4 h prior to analysis by fluorescence microscopy. Scale bar: 4 µm. **(D)** Quantification of the proportion of cells showing distinct foci in the experiment described in panel B. Bars indicate the mean of 1-3 biological replicates (diamonds). Number of cells analyzed in total: WT (1857), R93A (1818), R94A (1479), WT without *virS* (1477).

## Discussion

The identification of the ParB/Srx domain as a CTP-binding module has led to fundamental new insights into the function of ParAB*S* DNA partitioning systems. However, there are various ParB-like proteins that are not encoded in *parABS* operons and thus likely to mediate processes other than DNA segregation. Only very few representatives of these orphan ParB homologs have been investigated to date. One of them is the nucleoid occlusion protein Noc of *Bacillus subtilis*, a close homolog of chromosomally encoded ParB proteins that has recently been shown to use a CTP-dependent clamping-and-sliding mechanism to accumulate on chromosomal DNA and tether it to the cytoplasmic membrane^64^, thereby preventing the assembly of the cell division apparatus over the nucleoid^65, 66^. Another prominent member of this group is VirB, which has evolved into a transcriptional regulator with a critical role in *S. flexneri* virulence gene expression.

Structural modeling suggests that VirB is derived from plasmid-encoded ParB proteins and forms clamp-like dimeric structures that are closed by homodimerization of the NBDs and CTDs (**Figure 1**). In support of this model, previous work has shown that the two CTDs closely associate with each other and stably connect the two VirB subunits at their C-terminal ends^25^. Moreover, truncations of the NBD or CTD were found to completely abolish the function of VirB, underscoring the relevance of clamp formation and closure for its regulatory activity^24^. The NBD of VirB is highly conserved and binds CTP with similar or even higher affinity than previously characterized ParB homologs^38, 39^, with a clear preference for CTP over CDP (**Figure 2A,B**). Given the relatively high concentration of CTP in the cytoplasm (∼500 µM)^56^, the nucleotidebinding site of VirB is thus likely to be saturated with CTP at all times. Our biolayer interferometry, cross-linking and HDX analyses clearly demonstrate that CTP-binding facilitates the transition of VirB clamps from an open to a closed state in which they embrace target DNA in a ring-like fashion. As in the case of ParB, clamp closure by homodimerization of the CTP-bound NBDs leads to a crossover of the two polypeptide chains, positioning the NBD of one subunit next to the VBD of the respective *trans*-subunit (**Figure 1B**). However, these *trans*-interactions appear to be mediated by different interfaces and to be less extensive than those in the ParB dimer^36, 37, 39^. Importantly, we observe that CTP-mediated VirB clamp closure is stimulated by *virS* DNA (**Figures 6 and 7**). In low-stringency conditions, the addition of *virS* alone was sufficient to induce changes in the HDX pattern of the NBD that are indicative of transient NBD homodimerization events. This observation suggests that the juxtaposition of the VBDs at the *virS* site facilitates the face-to-face interaction of the two NBDs, likely by increasing their spatial proximity. The sole presence of CTP, by contrast, led to marked global changes in HDX that point to a more stable, but still transient association of the NBDs. Only if both *virS* and CTP were provided, VirB clamps closed robustly and with maximal efficiency. Notably, the crosslinking and HDX behavior of VirB in the presence of CTP is reminiscent of the behavior of ParB in the presence of CTPγS^37, 39^, consistent with the observation that VirB lacks significant CTPase activity under the conditions tested.

The strong non-specific DNA-binding activity of VirB complicates the analysis of its interaction with target promoters *in vitro*. In low-stringency buffer, VirB accumulated on DNA independently of the presence of CTP and *virS*, although DNA binding was more robust in the presence of CTP (**Figure 3**). Moreover, when analyzed in reactions containing CTP, non-specific DNA and *virS* DNA produced essentially the same changes in the HDX pattern of VirB (**Figures 7A and S9A**). This observation suggests that low-stringency conditions may allow *virS*-independent clamp closure, although it remains to be clarified whether this process occurs efficiently at cytoplasmic VirB and ligand concentrations. By contrast, in more stringent conditions that reduce the effect of non-specific protein-DNA interactions, VirB clamps are loaded specifically at *virS* sites, in a process strictly dependent on the presence of CTP (**Figure 4**). As observed for ParB, closed clamps are released from *virS* and slide laterally along the DNA, enabling the loading of multiple VirB dimers at a single *virS* site (**Figure 5**). Consistent with results previously obtained in *S. flexneri*^62^, our *in vivo* analysis confirmed that the presence of a single *virS* site is indeed sufficient to recruit a large fraction of VirB molecules to a *virS*-containing plasmid in *E. coli* cells (**Figure 8**). This process was abolished by the mutation of residues essential for CTP binding, verifying the critical relevance of CTP-dependent clamp closure for the association of VirB with target promoter regions. It still remains to be clarified how VirB can efficiently interact with *virS* sites despite the large excess of non-specific DNA within the cell. On the one hand, the lower salt concentration in the cytoplasm may be compensated by its high content of organic compounds and macromolecules, which could potentially also block the positively charged regions of VirB and, thus, reduce its non-specific DNA binding activity. On the other hand, the affinity of VirB for *virS* sites is likely to be markedly higher than that for non-specific DNA, since *virS* binding involves both non-specific interactions with the DNA backbone and specific interactions with the HTH-motif^25^. In support of this notion, previous work has shown that even a large (>50.000-fold) excess of non-specific DNA is not sufficient to prevent the formation of specific VirB-*virS* complexes^25^. Another open question concerns the mechanism that controls the dynamics of VirB clamp opening in the absence of appreciable CTPase activity. In the case of ParB, nucleotide hydrolysis was shown to serve two important purposes. On the one hand, it re-opens prematurely closed clamps to ensure the quantitative loading of ParB dimers onto DNA. On the other hand, it triggers robust clamp opening after loading and thus promotes the dissociation of VirB dimers from their target DNA, thereby determining the sliding time of ParB clamps and their degree of spreading within the centromere region^34, 36, 37^. However, at least in *Myxococcus xanthus*, CTPase-deficient ParB variants are still quantitatively loaded onto DNA, and they are still confined to a defined region within the centromere, although their longer sliding times lead to considerable increase in their spreading distances^37^. This observation is explained by spontaneous, CTPase-independent dissociation of the NBDs, which enables the release of ParB clamps without nucleotide hydrolysis, albeit at relatively low rates^34, 37^. VirB may rely on a similar mechanism to control its distribution within target promoter regions, and it will be interesting to study its spreading behavior and the kinetics of its release from DNA *in vivo*.

How can the loading and spreading of VirB clamps at *virS* sites affect promoter activity? VirB stimulates the expression of target genes by counteracting their silencing by H-NS^67^, a nucleoid-associated protein that can bridge DNA and thus stabilize negatively supercoiled DNA regions^68, 69^. Interestingly, the interaction of VirB with target promoter regions has recently been shown to generate torsional stress in target DNA molecules that induces the formation of positive supercoils^53^. In this way, it may remove adjacent negative supercoils and thus destabilize the H-NS nucleoprotein complexes that block transcription initiation at VirB-dependent promoters (**Figure 9**). In addition, the spreading of VirB clamps within the promoter region could sterically hinder the alignment and bridging of DNA by H-NS, thereby reinforcing this effect. The mechanism underlying the formation of positive supercoils by VirB clamps remains to be determined. It is conceivable that the two VBDs of a VirB dimers are bound slightly out of phase, so that their alignment upon clamp closure induces a small rotation of the associated *virS* half-sites in opposite directions, underwinding the DNA that is trapped within the clamp and slightly overwinding the flanking DNA regions. The strong non-specific DNA-binding activity may enable VirB to remain in close contact with the DNA after leaving *virS* and thus maintain this torsional force during the sliding process. The loading of many VirB clamps and their spreading within the promoter region may amplify the torsion generated and expand the size of the overwound region, thereby inducing extensive positive supercoiling. However, detailed structural studies are required to test this hypothesis.

**Figure 9.**
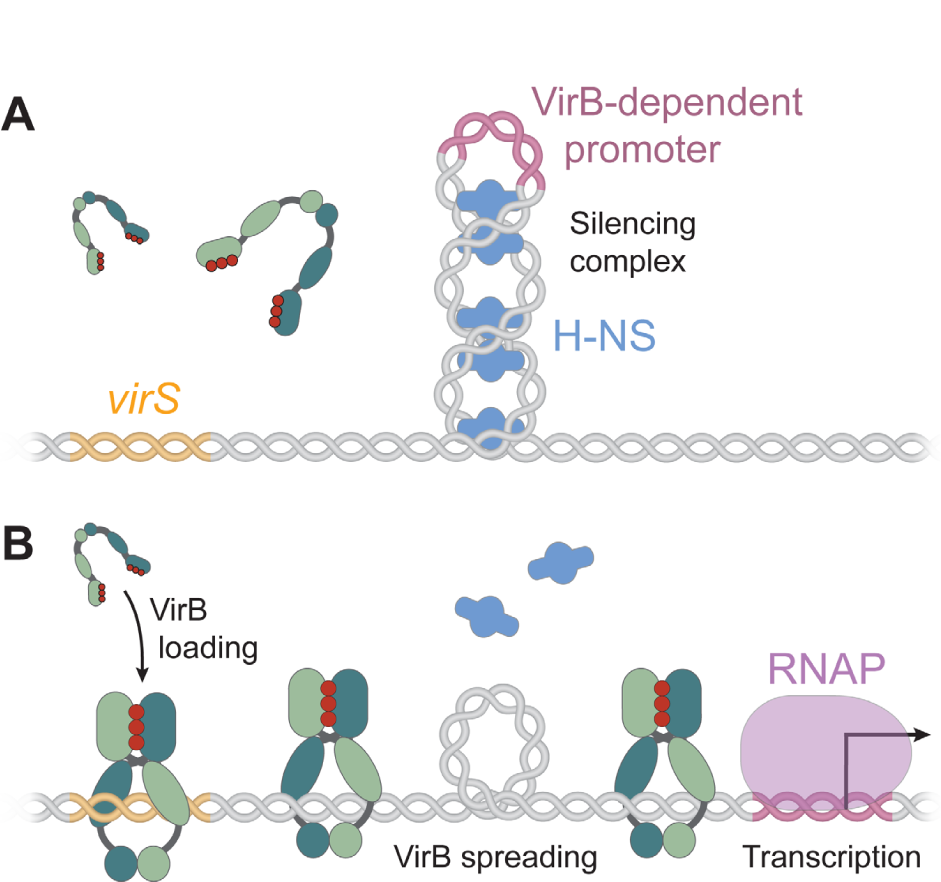
Hypothetical model of the mechanism underlying VirB-dependent gene regulation. **(A)** Before VirB associates with the virulence plasmid, the nucleoid-organizing protein H-NS binds and thus stabilizes negative DNA supercoils in the promoter regions of VirB-regulated genes, thereby sequestering the promoters from RNA polymerase and silencing gene expression. **(B)** The CTP-dependent loading of VirB clamps at *virS* sites and their spreading into the adjacent promoter regions leads to local overwinding of the DNA. This effect may destabilize the adjacent H-NS nucleoprotein complexes and reduce the degree of negative supercoiling in the vicinity of the promoter, thereby making it accessible to RNA polymerase and allowing transcription to occur.

Collectively, our work reveals that VirB forms a distinct group of ParB-like proteins that uses a CTP-dependent switch mechanism to associate with target promoter regions and activate gene expression. Future work will be required to fully unravel the connection between the loading and sliding of VirB clamps and their antagonistic effect on H-NS-mediated gene silencing. Given the diverse biological activities of orphan ParB homologs, it will be interesting to study more members of this intriguing group of proteins and determine the full breadth of functions they fulfill. Moreover, it is tempting to speculate that the distinctive nucleotide-binding domain and switch mechanism of VirB could be exploited for the development of antibacterial drugs that specifically suppress the induction of the *S. flexneri* virulence program.

## Methods

### Plasmids, strains and growth conditions

The plasmids and oligonucleotides used in this study are listed in **Tables S1-S2**. The sequences of all plasmids were verified by DNA sequencing. *E. coli* TOP10 (Invitrogen, USA) was used for cloning or localization studies. Proteins were overproduced in *E. coli* Rosetta(DE3)pLysS (Novagen, Germany). Cells were cultivated at 37 °C in Luria-Bertani (LB) broth supplemented with ampicillin (200 µg/mL) and chloramphenicol (34 µg/mL).

### Protein overproduction and purification

Wild-type VirB or mutant derivatives carrying an N-terminal His_6_-SUMO tag^70^ were overproduced in *E. coli* Rosetta(DE3)pLysS transformed with pSJ01, pSJ02, pSJ05, pSJ013 or pSJ014. Cultures were grown at 37 °C in 3 L LB medium with ampicillin and chloramphenicol to an OD_600_ of 0.6, induced to overproduce the protein of interest by the addition of 1 mM isopropyl-β-thiogalactopyranoside (IPTG), and further incubated at 18 °C overnight. The cells were harvested by centrifugation at 10,000 x g for 20 min at 4 °C and resuspended in lysis buffer (25 mM HEPES/NaOH pH 7.5, 500 mM NaCl, 0.1 mM EDTA, 5 mM MgCl_2_). After another centrifugation step at 6,500 x g for 30 min at 4 °C, the washed cells were resuspended in buffer A (25 mM HEPES/NaOH pH 7.5, 500 mM NaCl, 0.1 mM EDTA, 5 mM MgCl_2_, 30 mM imidazole, 1 mM β-mercaptoethanol) supplemented with 10 mg/mL DNase I and 100 µg/mL phenylmethylsulfonyl fluoride and disrupted by three passages through a French press (16,000 psi). The cell lysate was centrifuged at 30,000 x g for 30 min at 4 °C, and the supernatant was filtered through a syringe filter with a pore size of 0.2 μm. Subsequently, proteins were separated by immobilized-metal affinity chromatography (IMAC) on a 5 mL HisTrap HP column (GE Healthcare), previously washed with ddH_2_O and equilibrated with buffer B (25 mM HEPES/NaOH pH 7.5, 1 M NaCl, 0.1 mM EDTA, 5 mM MgCl_2_, 30 mM imidazole, 1 mM β-mercapto-ethanol). Protein was eluted at a flow rate of 1 mL/min with a linear gradient from 30 mM to 300 mM imidazole, obtained by mixing buffer B with buffer C (25 mM HEPES/NaOH pH 7.5, 500 mM NaCl, 0.1 mM EDTA, 5 mM MgCl_2_, 300 mM imidazole, 1 mM β-mercaptoethanol). Eluate fractions were analyzed by SDS-PAGE, and fractions containing the protein of interest in high concentration and purity were pooled and dialyzed overnight against 3 L of buffer D (25 mM HEPES/NaOH pH 7.5, 500 mM NaCl, 0.1 mM EDTA, 5 mM MgCl_2_, 10 % (v/v) glycerol, 1 mM β-mercaptoethanol). After the removal of precipitates by centrifugation at 30,000 x g for 30 min at 4 °C, the protein solution was filtered as described above. Subsequently, the His_6_-SUMO tag was removed by treatment with Ulp1 protease^70^ in the presence of 1 mM DTT, while the protein solution was dialyzed overnight against buffer D. A second IMAC step was then performed to separate the cleaved His_6_-SUMO tag from the protein of interest. Suitable flow-through fractions were pooled and dialyzed overnight against buffer D. After concentration to a final volume of 5 mL, the solution was applied to a size exclusion chromatography (SEC) on a HighLoad 75 Superdex column (GE Healthcare) equilibrated with buffer D. Fractions containing pure protein of interest were pooled, concentrated, snap-frozen in liquid N_2_ and stored at −80 °C until further use. *M. xanthus* ParB was purified as described previously^37^.

### Isothermal titration calorimetry

Nucleotide binding assays using isothermal titration calorimetry were performed with a MicroCal PEAQ-ITC system (Malvern Panalytical, USA) at 25 °C. Prior to the measurements, CTPγS (custom synthesized by Jena Biosciences, Germany) and CDP were dissolved to a concentration of 1.55 mM in reaction buffer (25 mM HEPES/NaOH pH 8, 150 mM NaCl, 0.1 mM EDTA, 5 mM MgCl_2_). Subsequently, the nucleotide solutions were titrated to 115 μM VirB in 13 consecutive injections (2 μL), performed at 150-s intervals with a duration of 4 s per injection. The mean enthalpies of dilution were subtracted from the raw titration data before analysis. The titration curves obtained were fitted to a one-set-of-sites model using the MicroCal PAEQ-ITC analysis software (Malvern Panalytical).

### Microscale thermophoresis

Nucleotide binding assays based on microscale thermophoresis were performed with a Monolith NT.115 instrument (NanoTemper Technologies GmbH, Germany) using Monolith NT Premium Capillaries. Proteins were fluorescently labeled using the RED-Maleimide 2nd Generation Protein Labeling Kit (NanoTemper Technologies GmbH, Germany) as recommended by the manufacturer. 50 nM labeled protein was then mixed with CTP, CDP or ATP at concentrations ranging from 61 nM to 2 mM in MST buffer (25 mM HEPES/NaOH pH 7.2, 500 mM NaCl, 5 mM MgCl_2_, 7.5% (v/v) glycerol, 0.06% [v/v] Tween-20). Measurements were performed at 25 °C, with the red LED laser adjusted to a power of 50 % (VirB-R93A) or 70 % (VirB and VirB-R94A) and the infrared laser set to 50 %. Two to three independent measurement (three technical replicates each) were performed for each condition. Data were analyzed using MO Affinity Analysis v2.3 (NanoTemper Technologies GmbH, Germany). To prevent heat-induced artifacts and, at the same time, avoid analyzing only the signal change induced by the initial temperature jump, the following regions were used for data analysis: cold region from −1 s to 0 s, hot region from 1.5 sec to 2.5 sec.

### CTPase assay

Nucleotide hydrolysis was analyzed using an NADH-coupled enzyme assay ^71, 72^. The reactions contained 1 mM CTP, 5 μM protein (VirB or ParB), 800 μg/mL NADH, 20 U/mL L-lactate dehydrogenase (Sigma Aldrich), 20 U/mL pyruvate kinase (Sigma Aldrich) and 3 mM PEP in a final volume of 200 µl. The reaction buffer used for VirB comprised 25 mM HEPES/NaOH pH 7.5, 5 mM MgCl_2_, 500 mM NaCl, 0.1 mM EDTA and 1 mM DTT and included 0.5 µM of a double-stranded DNA oligonucleotide (23 bp) with a *virS* site (virS-icsB-for/virS-icsB-rev) or a scrambled *virS* site (Scrambled-virS-for/Scrambled-virS-rev) (**Table S3**). For ParB, the reaction buffer comprised 25 mM HEPES/NaOH pH 7.2, 5 mM MgCl_2_, 150 mM NaCl and 1 mM DTT and included 0.3 µM of a DNA stem-loop (54 bases; *parS*-Mxan-wt) with a wild-type *parS* site (**Table S3**). After the start of the reactions by the addition of CTP, 150 μL of each mixture were transferred to a 96-well plate (Sarstedt, Germany). Subsequently, the absorbance of NADH at a wavelength of 340 nm was measured over 45 min at 2-min intervals in an Epoch 2 microplate spectrometer (BioTek, Germany), which was set to a temperature of 30 °C. A control reaction lacking protein was analyzed to correct for NADH oxidation and/or spontaneous CTP hydrolysis. Data were analyzed and plotted using Excel 2019 (Microsoft). The data were fitted to a linear equation, whose slope was then used to calculate the turnover numbers of VirB and ParB for CTP.

### Biolayer interferometry

Biolayer interferometry (BLI) analyses were performed using a Bli(tz) system (ForteBio, Pall Life Science), equipped with High Precision Streptavidin (Octet SAX2) biosensors (Sartorius, USA) that were derivatized with biotinylated DNA fragments, Short, single-biotinylated DNA fragments (23 bp) were assembled from two oligonucleotides (Eurofins, Germany), which were mixed, heated to 95 °C for 5 min and then annealed by gradual reduction of the temperature. Long, double-biotinylated DNA fragments (215 bp) were obtained by PCR amplification of custom-synthesized DNA fragments (Strings^TM^ DNA Fragments; Invitrogen, Germany) with biotinylated primers (Bio-icsB-for/Bio-icsB-rev), followed by gel purification. Reactions were carried out in low-stringency (25 mM HEPES/NaOH pH 7.5, 150 mM NaCl, 5 mM MgCl_2_, 1 mM DTT, 10 μM BSA, 0.01% [v/v] Tween 20) or high-stringency (25 mM HEPES/NaOH pH 7.5, 500 mM NaCl, 5 mM MgCl_2_, 1 mM DTT, 10 μM BSA, 0.01% [v/v] Tween 20) binding buffer. After the establishment of a stable baseline, the association reactions were monitored at different concentrations of VirB in binding buffer. To monitor the dissociation kinetics, the sensor was subsequently transferred to a protein-free buffer. The data obtained were analyzed using Excel 2019 (Microsoft).

To analyze VirB for sliding behavior, the closed 215-bp *virS*-containing DNA fragment was open by cleavage with NdeI (NEB, Germany). To this end, biosensors with immobilized DNA were incubated overnight at 37°C with 80 U *Nde*I in 300 µL rCutSmart buffer (NEB, Germany). In parallel, a control biosensor was prepared by incubation in 300 µL rCutSmart buffer in the absence of NdeI. Subsequently, the biosensors were probed with VirB in high-stringency buffer as described above.

### Hydrogen-deuterium exchange (HDX) mass spectrometry

HDX-MS experiments were carried out similarly as described previously^37, 38^ with minor modifications. In HDX experiment 1 (low-stringency buffer), the samples contained 25 µM VirB in a buffer composed of 25 mM HEPES-NaOH pH 7.5, 150 mM NaCl, 5 mM MgCl_2_, 0.1 mM EDTA and 1 mM DTT. In HDX experiment 2 (high-stringency buffer), the samples contained 50 µM VirB in a buffer composed of 25 mM HEPES-NaOH pH 7.5, 500 mM NaCl, 5 mM MgCl_2_, 1 mM DTT and 0.1 mM EDTA. When indicated, double-stranded DNA oligonucleotides containing a native or scrambled *virS* sequence were used at the same concentration as VirB, and CTP was used at a final concentration of 10 mM.

HDX reactions were prepared automatically with a two-arm robotic autosampler (LEAP Technologies). 7.5 µL of protein solution was dispensed in a glass well plate kept at 25 °C and supplemented with 67.5 μL of buffer (see above) prepared in 99.9 % D_2_O to initiate the hydrogen/deuterium exchange reaction. After incubation for 10, 30, 100, 1,000 or 10,000 s, 55 μL of the reaction were withdrawn and added to 55 µL quench buffer (400 mM KH_2_PO_4_/H_3_PO_4_, 2 M guanidine-HCl, pH 2.2), which was pre-dispensed in another glass well plate cooled at 1 °C. Following mixing, 95 µL of the resulting mixture were injected into an ACQUITY UPLC M-Class System with HDX Technology through a 50 µL sample loop. Non-deuterated samples were prepared using a similar procedure by 10-fold dilution in buffer prepared with H_2_O followed by an ∼10 s incubation at 25 °C prior to quenching and injection. The samples were flushed out of the sample loop with water + 0.1 % (v/v) formic acid (100 µL/min flow rate) over 3 min and guided to a cartridge (2 mm x 2 cm) that contained immobilized porcine pepsin for proteolytic digestion at 12 °C. The resulting peptic peptides were collected on a trap column cartridge (2 mm x 2 cm) that was filled with POROS 20 R2 reversed-phase resin (Thermo Scientific) and kept at 0.5 °C for 3 min, after which the trap column was placed in line with an ACQUITY UPLC BEH C18 1.7 μm 1.0 x 100 mm column (Waters)^73^. Peptides were eluted at 0.5 °C with a gradient of water + 0.1% (v/v) formic acid (eluent A) and acetonitrile + 0.1% (v/v) formic acid (eluent B) at 60 µL/min flow rate as follows: 0-7 min/95-65% A, 7-8 min/65-15% A, 8-10 min/15% A, guided to a G2-Si HDMS mass spectrometer with ion mobility separation (Waters) and ionized by electrospray ionization (capillary temperature 250 °C, spray voltage 3.0 kV). Mass spectra were acquired over a range of 50 to 2,000 m/z in enhanced high-definition MS (HDMS^E^)^74, 75^ or high-definition MS (HDMS) mode for non-deuterated and deuterated samples, respectively. [Glu1]-Fibrinopeptide B standard (Waters) was employed for lock mass correction. During separation of the peptides, the pepsin column was washed three times with 80 µL of 4% (v/v) acetonitrile and 0.5 M guanidine hydrochloride, and blank runs (injection of H_2_O) were performed between each sample. Three technical replicates (independent H/D exchange reactions) were measured per incubation time. No correction for HDX back exchange was conducted.

Further data analysis was conducted as described previously^37, 38^. Peptides were identified with ProteinLynx Global SERVER 3.0.1 (PLGS, Waters) from the non-deuterated samples acquired with HDMS^E^ by employing low energy, elevated energy, and intensity thresholds of 300, 100 and 1,000 counts, respectively. Hereby, the identified ions were matched to peptides with a database containing the amino acid sequences of VirB, porcine pepsin and their reversed sequences with the following search parameters: peptide tolerance = automatic; fragment tolerance = automatic; min fragment ion matches per peptide = 1; min fragment ion matches per protein = 7; min peptide matches per protein = 3; maximum hits to return = 20; maximum protein mass = 250,000; primary digest reagent = non-specific; missed cleavages = 0; false discovery rate = 100. Deuterium incorporation into peptides was quantified with DynamX 3.0 software (Waters). Only peptides that were identified in all non-deuterated samples and with a minimum intensity of 10,000 counts, a maximum length of 40 amino acids, a minimum number of two products, a maximum mass error of 25 ppm, and retention time tolerance of 0.5 minutes were considered for analysis. All spectra were manually inspected and, if necessary, peptides were omitted (e.g., in case of low signal-to-noise ratio or presence of overlapping peptides). Mass spectra of VirB samples containing scrambled *virS* were generally of lower quality than the other states and could only be partially assigned.

The observable maximal deuterium uptake of a peptide was calculated by the number of residues minus one (for the N-terminal residue, which after proteolytic cleavage quantitatively loses its deuterium label) minus the number of proline residues contained in the peptide (which lack an exchangeable peptide bond amide proton). For the calculation of HDX in percent, the absolute HDX was divided by the theoretical maximal deuterium uptake and multiplied by 100. The rendering of residue-specific HDX differences from overlapping peptides for any given residue of VirB was performed with DynamX 3.0 by employing the shortest peptide covering any residue. Where multiple peptides were of the shortest length, the peptide with the residue closest to the C-terminus of the peptide was used.

### *In vitro* crosslinking

Prior to the crosslinking reactions, VirB or its mutant variants were incubated with 5 mM Tris(2-carboxyethyl) phosphine (TCEP; Sigma, USA) for 1 h at room temperature to fully reduce all cysteine residues. Subsequently, the protein was transferred to reaction buffer (25 mM HEPES/NaOH pH 7.5, 500 mM NaCl, 5 mM MgCl_2_, 0.1 mM EDTA) using PD SpinTrap G-25 columns (Cytiva, Germany). Crosslinking was performed at room temperature in mixtures containing 10 µM protein, which were supplemented with 1 mM CTP and/or 1 µM of a *virS*-containing double-stranded DNA oligonucleotide (virS-icsB-for/virS-icsB-rev) when appropriate. After the indicated incubation times, bismaleimidoethane (BMOE; dissolved in dimethylsulfoxide) was added to a concentration of 1 mM. After brief mixing and incubation for 1 min, the reaction was stopped by the addition of dithiothreitol-containing SDS sample buffer. As a negative control, samples were treated with dimethylsulfoxide instead of BMOE. The samples were then analyzed by SDS-polyacrylamide gel electrophoresis, and protein was stained with InstantBlue Coomassie Protein Stain (Expedeon, Germany). The gels were imaged with a ChemiDoc MP imaging system (Bio-Rad Laboratories, USA), and the intensities of the different bands were quantified using Image Lab software (Bio-Rad Laboratories, USA).

### Nucleotide content analysis

10 µL of VirB (50 µM) were mixed with 40 µL double-distilled water and 150 µL chloroform, followed by 5 s of vigorous shaking, 15 s of heat denaturation at 95 °C and snap-freezing in liquid nitrogen. After thawing and centrifugation (17,300 x g, 10 min, 4 °C), the aqueous phase containing any released nucleotides was withdrawn for HPLC analysis on an Agilent 1260 Infinity system equipped with a Metrosep A Supp 5 - 150/4.0 column (Metrohm). Samples (10 µL) were injected and eluted at a flow rate of 0.6 mL/min flow rate with 90 mM (NH_4_)_2_CO_3_ pH 9.25. Nucleotides were detected at 260 nm wavelength by comparison of their retention time with those of a commercial standard that was treated as described for the VirB samples.

### Fluorescence microscopy

Cells were inoculated to an OD_600_ 0.03 in LB medium and grown for 1 h at 28°C before the medium was supplemented with 0.1 % (w/v) arabinose to induce the production of the indicated mVenus-VirB variants. After further cultivation to an OD_600_ of 0.6, cells were immobilized on 1 % (w/v) agarose pads and imaged with a Zeiss Axio Observer.Z1 microscope (Zeiss, Germany) equipped with a Zeiss Plan-Apochromat 100x/1.40 Oil Ph3 M27 objective and a pco.edge 4.2 sCMOS camera (PCO, Germany). An X-Cite 120PC metal halide light source (EXFO, Canada) and ET-YFP filter cubes (Chroma, USA) were used for fluorescence detection. Images were recorded with VisiView 3.3.0.6 (Visitron Systems, Germany) and processed with Fiji ^76^ and Adobe Illustrator CS6 (Adobe Inc., San Jose, USA). All microscopic analyses were performed in triplicate, using independent biological replicates.

## Acknowledgements

We thank Olga Ebers for excellent technical assistance, Sven Freibert for advice on microscale thermophoresis, Andreas Diepold for providing pCP301, and Lucas Schnabel for helpful discussions. This work was supported by the University of Marburg (core funding to M.T.), the Max Planck Society (Max Planck Fellowships to G.B. and M.T.) and the German Research Foundation (DFG) (project 269423233 – TRR 174 to M.T. and project 324652314 – DFG Core Facility for Interactions, Dynamics and Biomolecular Assembly Structure to G.B.).

## Data availability

All data generated in this study are included in the manuscript and the supplemental material.

## Author contributions

S.J. constructed plasmids, purified proteins and performed biochemical and fluorescence microscopy studies. W.S. conducted the hydrogen-deuterium exchange mass spectrometric analyses. J.H. performed the microscale thermophoresis analysis. J.R. purified constructed plasmids, purified proteins and conduced fluorescence microscopy studies. P.I.G. performed the isothermal calorimetry analysis. S.J., W.S., J.H., M.O.V., P.I.G. and M.T. analyzed the data. G.B. and M.T. secured funding and supervised the study. M.T. conceived the study. S.J. and M.T. wrote the paper, with input from all other authors.

## Competing interests

The authors declare no competing interests.

## Supplementary figures

**Figure S1.**
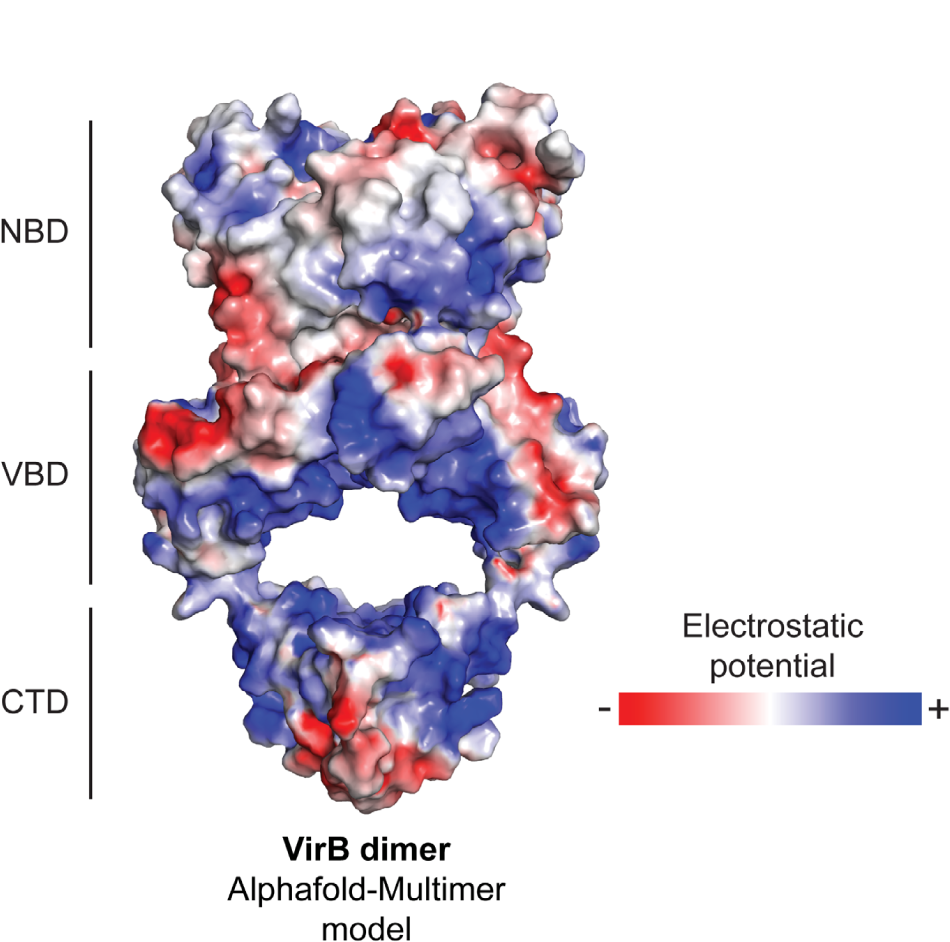
Electrostatic potential surface of VirB. The electrostatic potential surface of VirB was determined with PyMOL (Schrödinger LLC, USA) and plotted onto a structural model of the VirB dimer, generated with AlphaFold-Multimer (Evans et al, 2022). The nucleotide-binding domain (NBD), the *virS*-binding domain (VBD) and the C-terminal dimerization domain (CTD) are indicated. The color code is given on the right.

**Figure S2.**
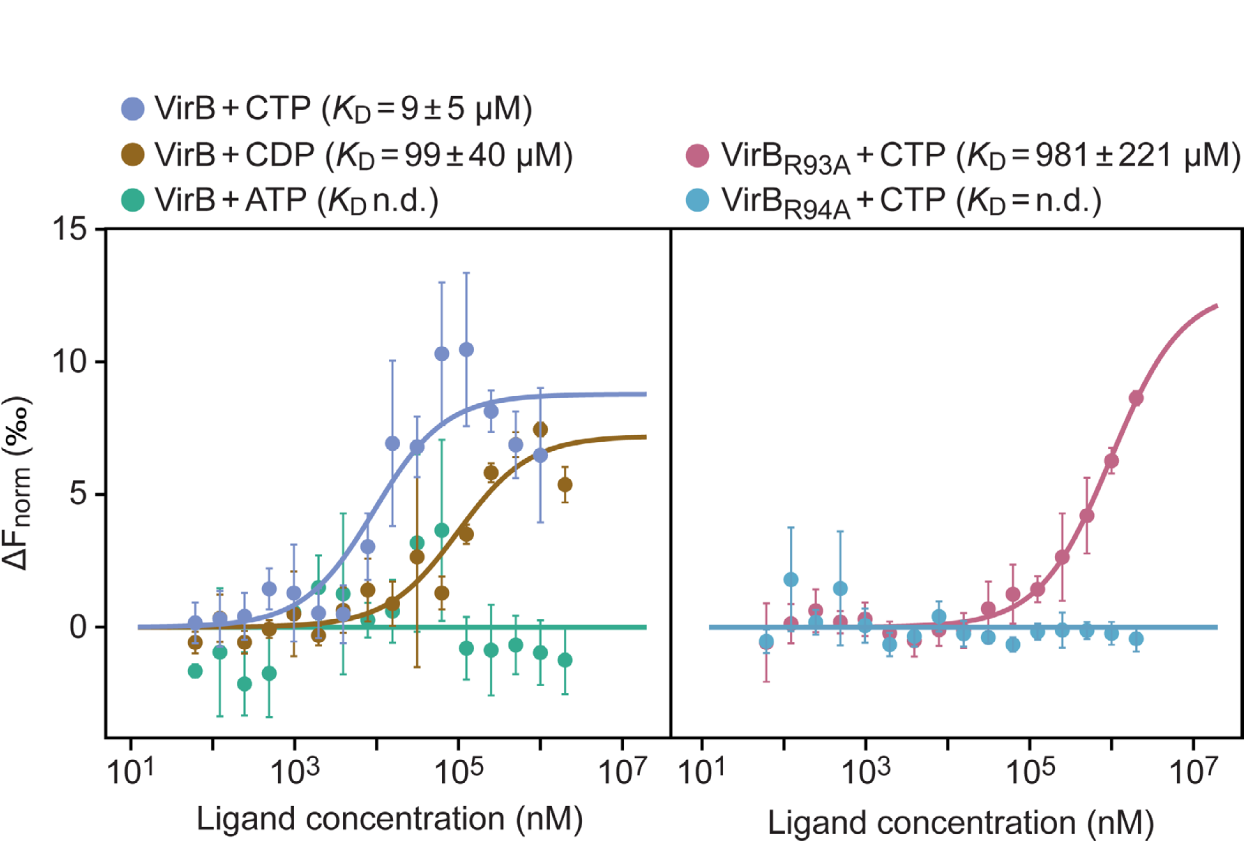
Microscale thermophoresis analysis of the nucleotide-binding behavior of VirB and mutant variants. Wild-type VirB, VirB-R93A and VirB-R94A were fluorescently labeled and then analyzed for their thermophoretic mobility in the presence of increasing concentrations of nucleotides. The data points represent the mean of the normalized ΔF values (± SD) obtained in 2-3 independent experiments (each performed in triplicate). To determine the binding affinities, the results of all replicates were combined and fitted using MO.Affinity Analysis v2.3 (Nanotemper, Germany). The *K*_D_ values obtained are indicated above the graphs. n.d. = not detectable.

**Figure S3.**
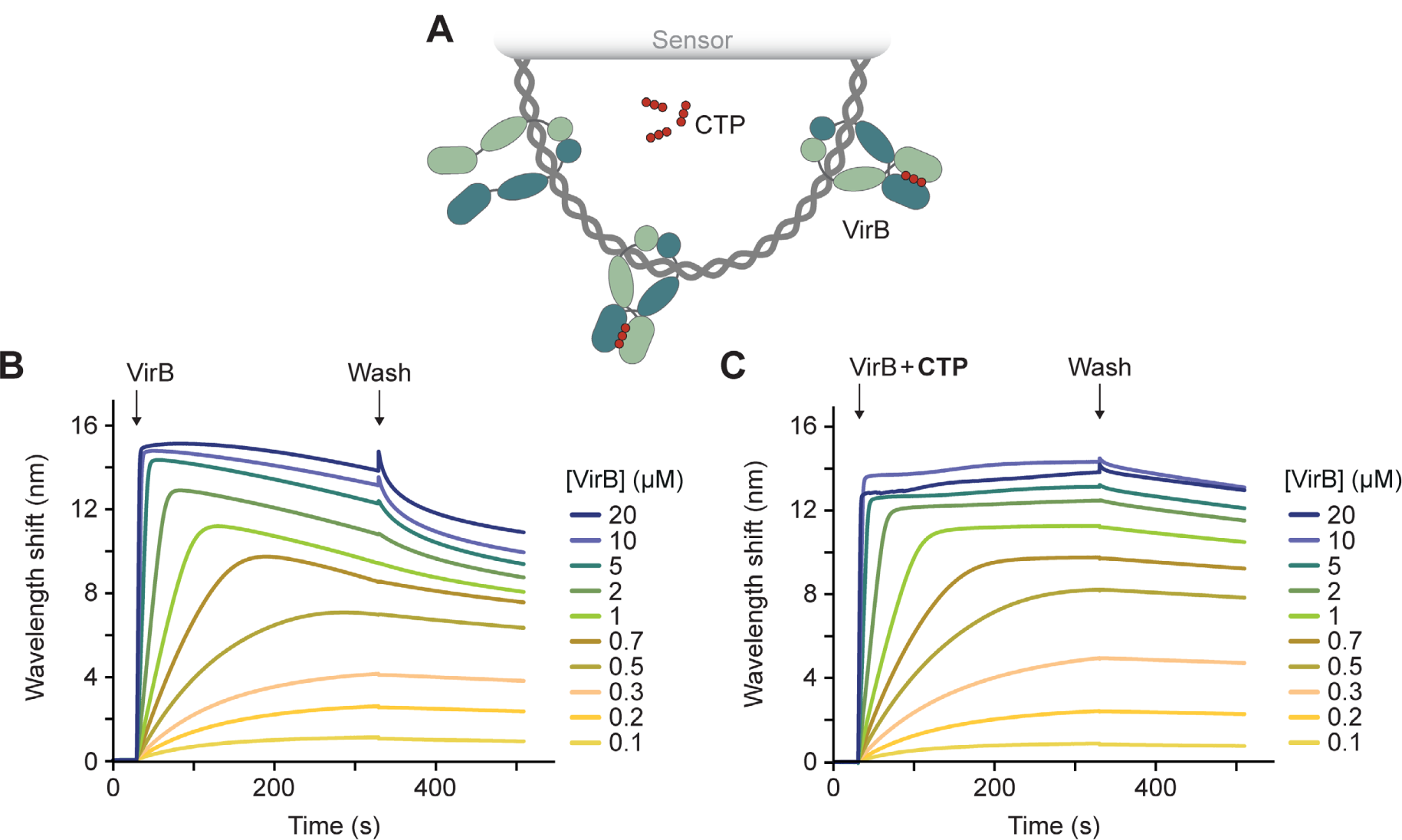
Biolayer interferometry analysis of the interaction of VirB with closed non-specific DNA in low-stringency conditions. **(A)** Schematic of the biolayer interferometry setup used for the analyses in panels B and C. A double-biotinylated dsDNA fragment (215 bp) containing a scrambled *virS* sequence was immobilized on a streptavidin-coated biosensor and probed with VirB (green). **(B,C)** BLI analysis of the DNA-binding behavior of VirB in (B) the absence and (C) the presence of CTP (1 mM) in low-stringency buffer (150 mM NaCl). Sensors carrying the closed DNA fragment (at a density corresponding to a wavelength shift of ∼1.3 nm) were probed with the indicated concentrations of VirB. At the end of the association reactions, the biosensors were transferred into protein- and nucleotide-free buffer to monitor the dissociation reactions (wash). The graphs show the results of a representative experiment (n=2-3 independent replicates).

**Figure S4.**
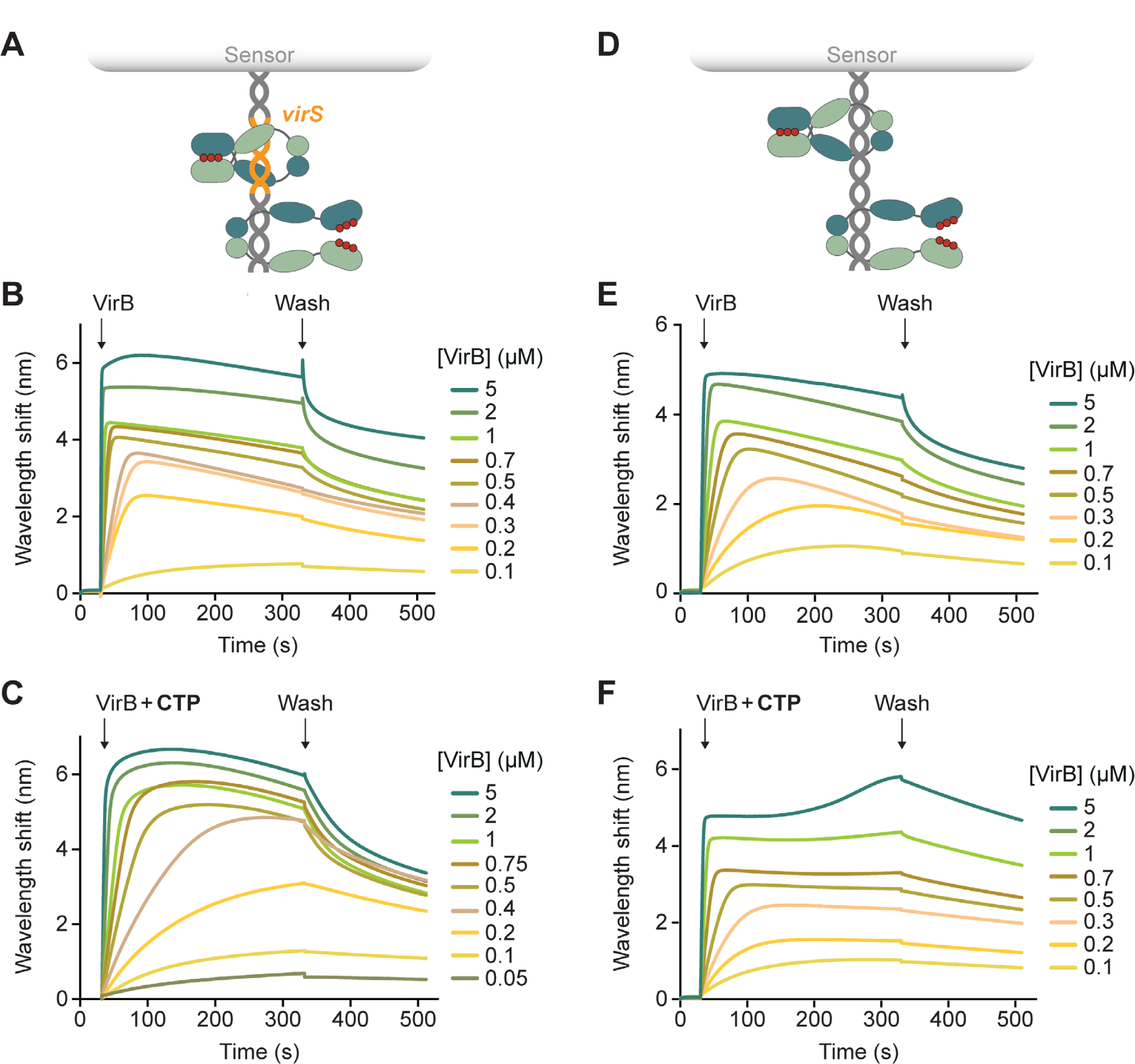
Biolayer interferometry analysis of the interaction of VirB with an open double-stranded oligonucleotide in low-stringency conditions. **(A)** Schematic of the biolayer interferometry setup used for the analysis in panels B and C. A double-stranded *virS*-containing DNA oligonucleotide biotinylated at one of its ends was immobilized on a streptavidin-coated biosensor. **(B,C)** DNA-binding behavior of VirB in the (B) absence and (C) presence of CTP (1 mM) in low-stringency buffer (150 mM NaCl). Biosensors carrying the open *virS*-containing target DNA (at a density corresponding to a wavelength shift of ∼1.5 nm) were probed with the indicated concentrations of VirB. The graphs show the results of a representative experiment (n=2-3 independent replicates). **(D)** Schematic of the biolayer interferometry (BLI) setup used for the analysis in panels E and F. A double-stranded DNA oligonucleotide containing a scrambled *virS* sequence and carrying a biotin moiety at one of its ends was immobilized on a streptavidin-coated biosensor. **(E,F)** The open, non-specific target DNA was probed with VirB in the (E) absence and (F) presence of CTP (1 mM) as described for panels B and C.

**Figure S5.**
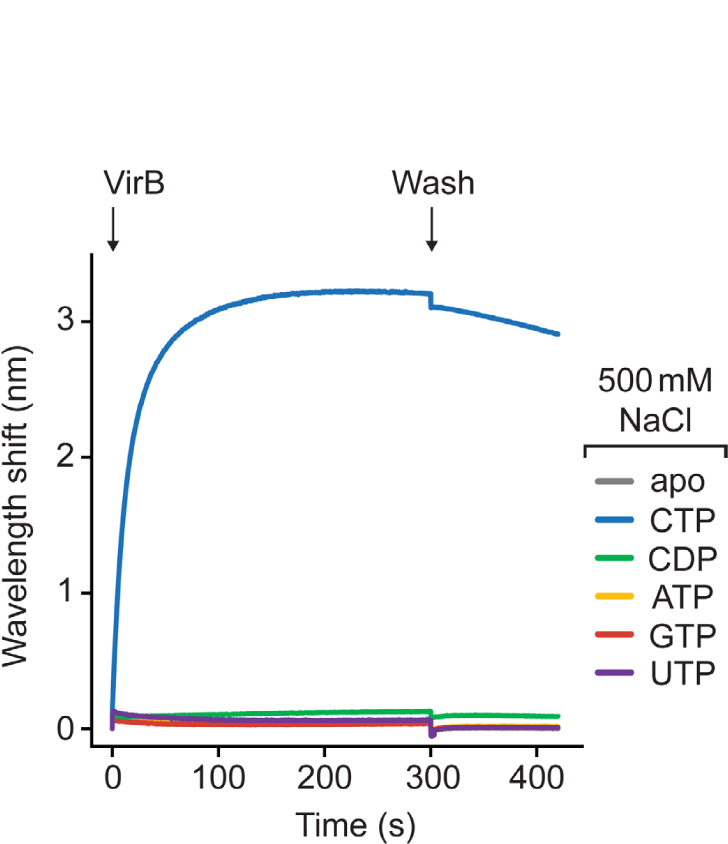
Biolayer interferometry analysis of the interaction of VirB with closed, *virS*-containing DNA in the presence of different nucleotides. A streptavidin-coated biosensor carrying a double-biotinylated *virS*-containing DNA fragment (215 bp) was probed with VirB (20 µM) in the presence of the indicated nucleotides (1 mM) in high-stringency buffer (500 mM NaCl).

**Figure S6.**
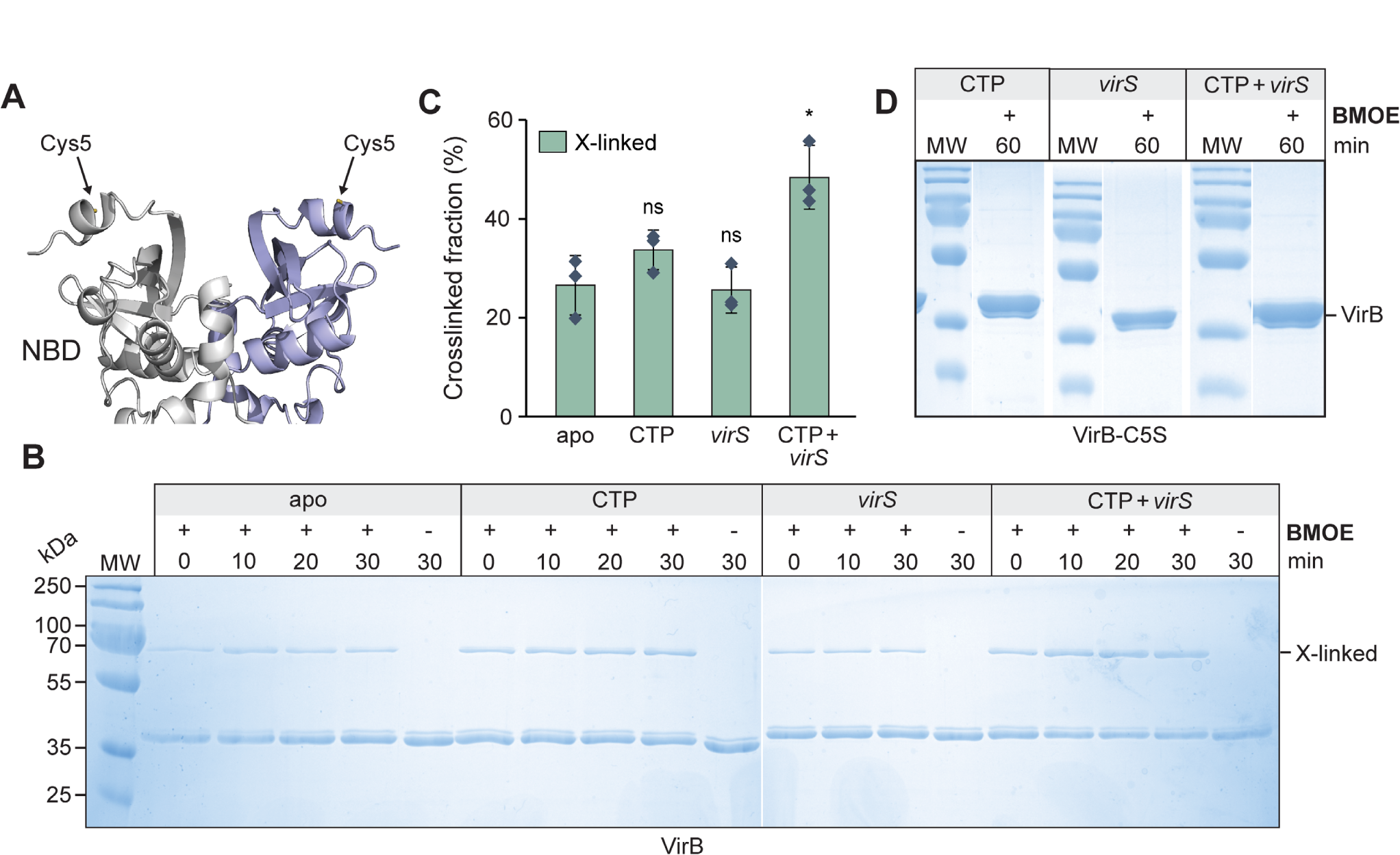
*In vitro* crosslinking analysis of wild-type VirB. **(A)** Close-up view of the N-terminal region of the VirB dimer, as modeled by AlphaFold-Multimer ^1^. Arrow indicate the predicted positions of the native cysteine residues (C5). **(B)** SDS-polyacrylamide gels showing the protein species obtained in the *in vitro* crosslinking analysis. Wild-type VirB (10 µM) was incubated for the indicated time periods with CTP (1 mM), with a double-stranded DNA oligonucleotide containing a *virS* sequence (1 µM; virS-icsB-for/virS-icsB-rev) or with both CTP and *virS* DNA prior to crosslinking with BMOE and analysis of the reaction products by SDS-PAGE. Monomeric VirB and the dimeric crosslinking product (X-linked) are indicated. MW: Molecular weight marker. **(C)** Quantification of the fractions of crosslinked protein obtained in the indicated conditions. The columns show the mean (± SD) of three independent measurements (diamonds). *p<0.05, ns: not significant (Welch’s t-test; versus the apo state). **(D)** SDS-gel showing the results of control *in vitro* crosslinking analyses performed with the cysteine-free VirB-C5S variant. The reactions were performed as described in panel B, with an incubation time of 60 min prior to crosslinking and SDS-PAGE analysis.

**Figure S7.**
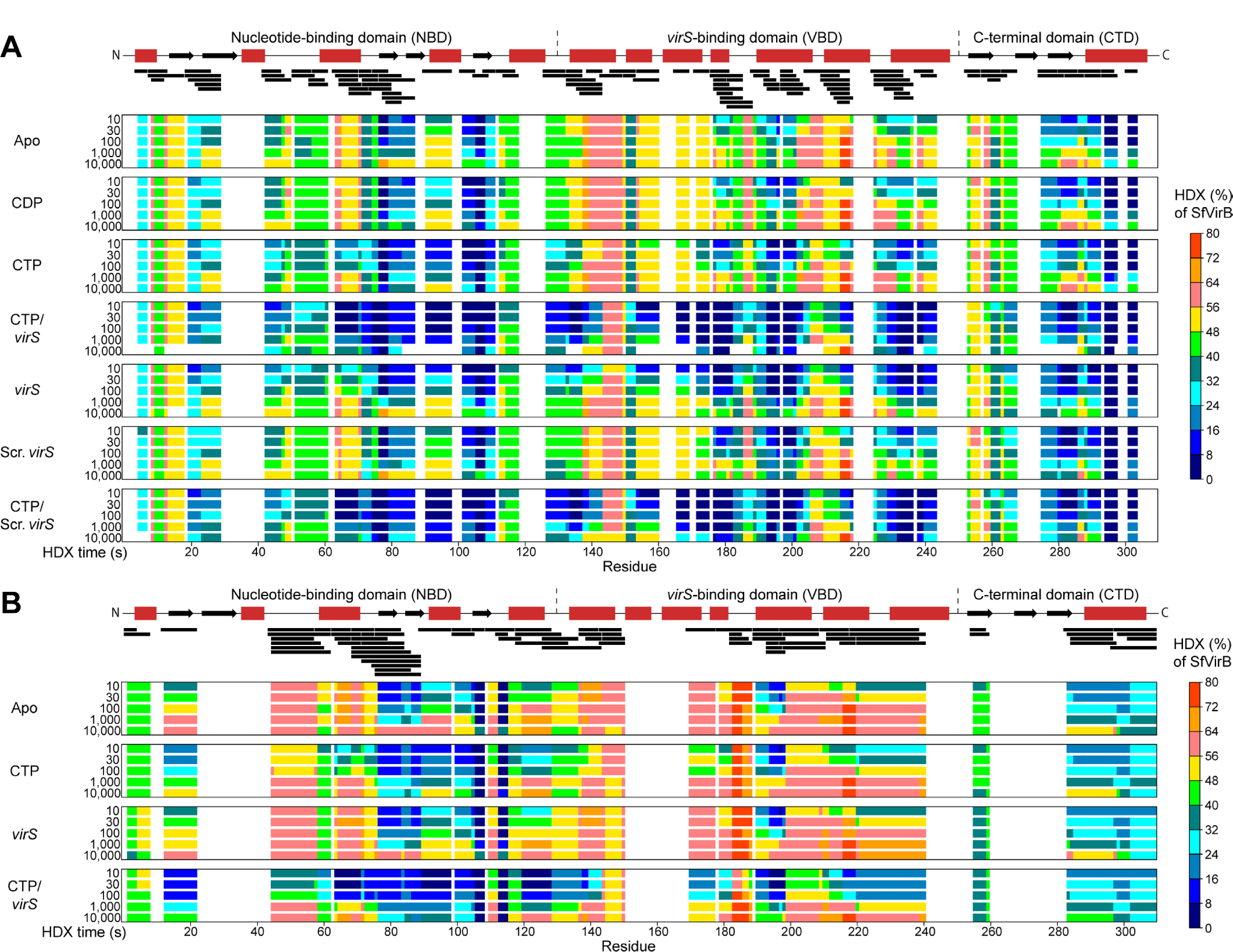
Hydrogen-deuterium exchange (HDX) analysis of VirB. **(A)** HDX analysis of VirB in low-stringency conditions (150 mM NaCl). VirB (25 µM) was incubated in deuterated buffer for the indicated time intervals alone (apo), with double-stranded DNA oligonucleotides containing a scrambled (scrambled-virS-for/scrambled-virS-rev) or intact (virS-icsB-for/virS-icsB-rev) *virS* motif (25 µM) and/or with the indicated nucleotides (10 mM) prior to HDX analysis. Shown is the degree of HDX along the primary sequence of VirB in the indicated conditions. The color scale is given on the right. The schematic at the top displays the predicted secondary structure of VirB. The black bars represent peptides of VirB that could be analyzed for HDX. Residue-specific HDX information was obtained from these overlapping peptides by employing the shortest peptide covering any residue. No HDX could be obtained for amino acid sequences in the gaps, which indicate regions not covered by any peptides. **(B)** HDX analysis of VirB in high-stringency conditions (500 mM NaCl). VirB (50 µM) was incubated in deuterated buffer for the indicated time intervals alone (apo), with double-stranded DNA oligonucleotides containing a scrambled or intact *virS* motif (50 µM) and/or CTP (10 mM) prior to HDX analysis. The data are presented as described for panel A. Detailed information about the peptides analyzed to generate the graphs in panels A and B is given in **Data S1**.

**Figure S8.**
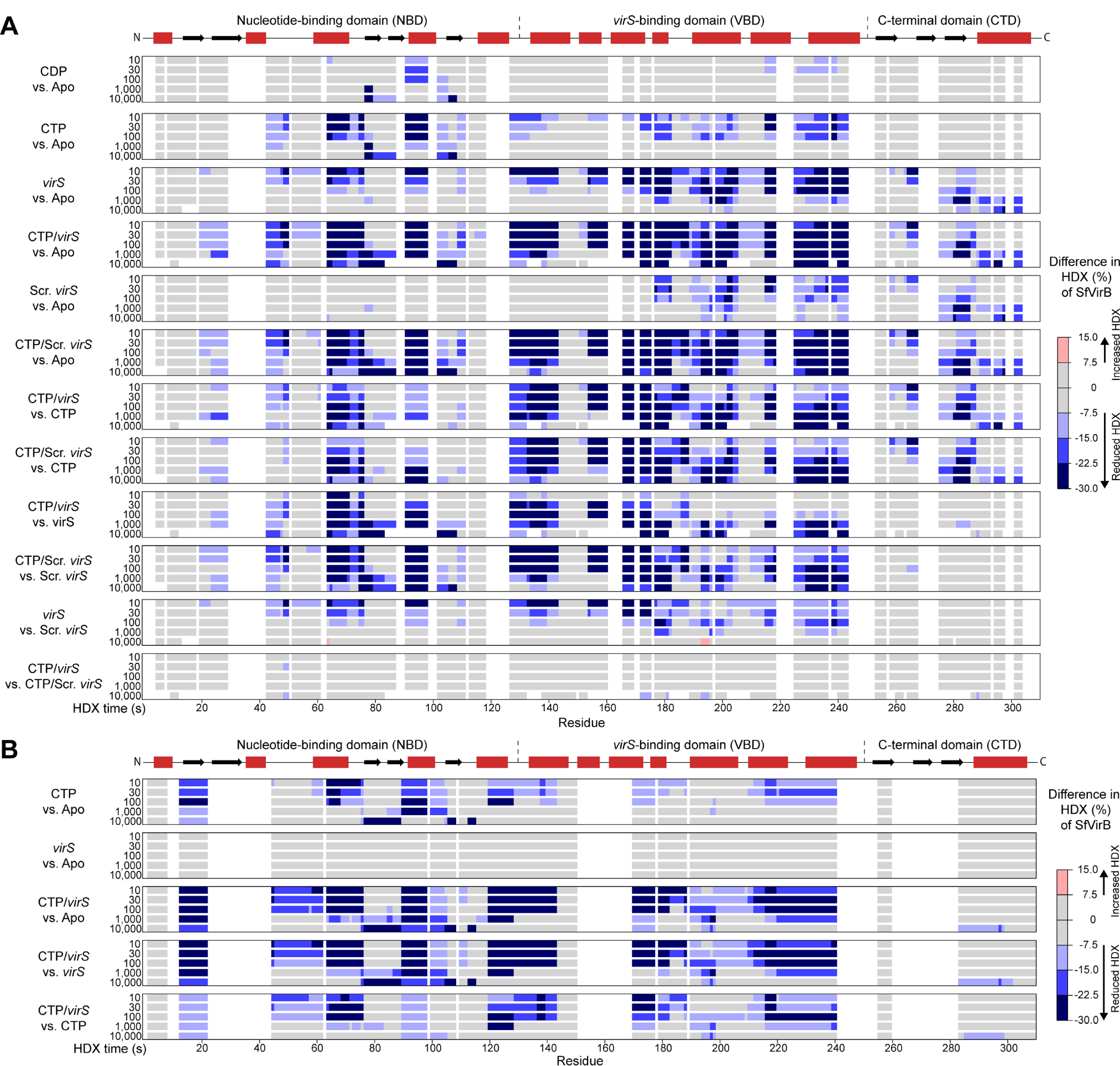
Ligand-induced changes in the HDX pattern of VirB. **(A)** HDX analysis of VirB in low-stringency conditions (150 mM NaCl). Shown are residue-specific differences in HDX between the indicated states of VirB, projected onto the amino acid sequence of VirB. The results are derived from pairwise comparisons of the HDX data presented in **Figure S7**. The color code is given on the right. Blue color denotes reduced HDX in the first state compared to the second state in “first state” vs. “second state” comparisons. The schematic at the top shows the predicted secondary structure of VirB. **(B)** HDX analysis of VirB in high-stringency conditions (500 mM NaCl). The data are presented as described for panel A.

**Figure S9.**
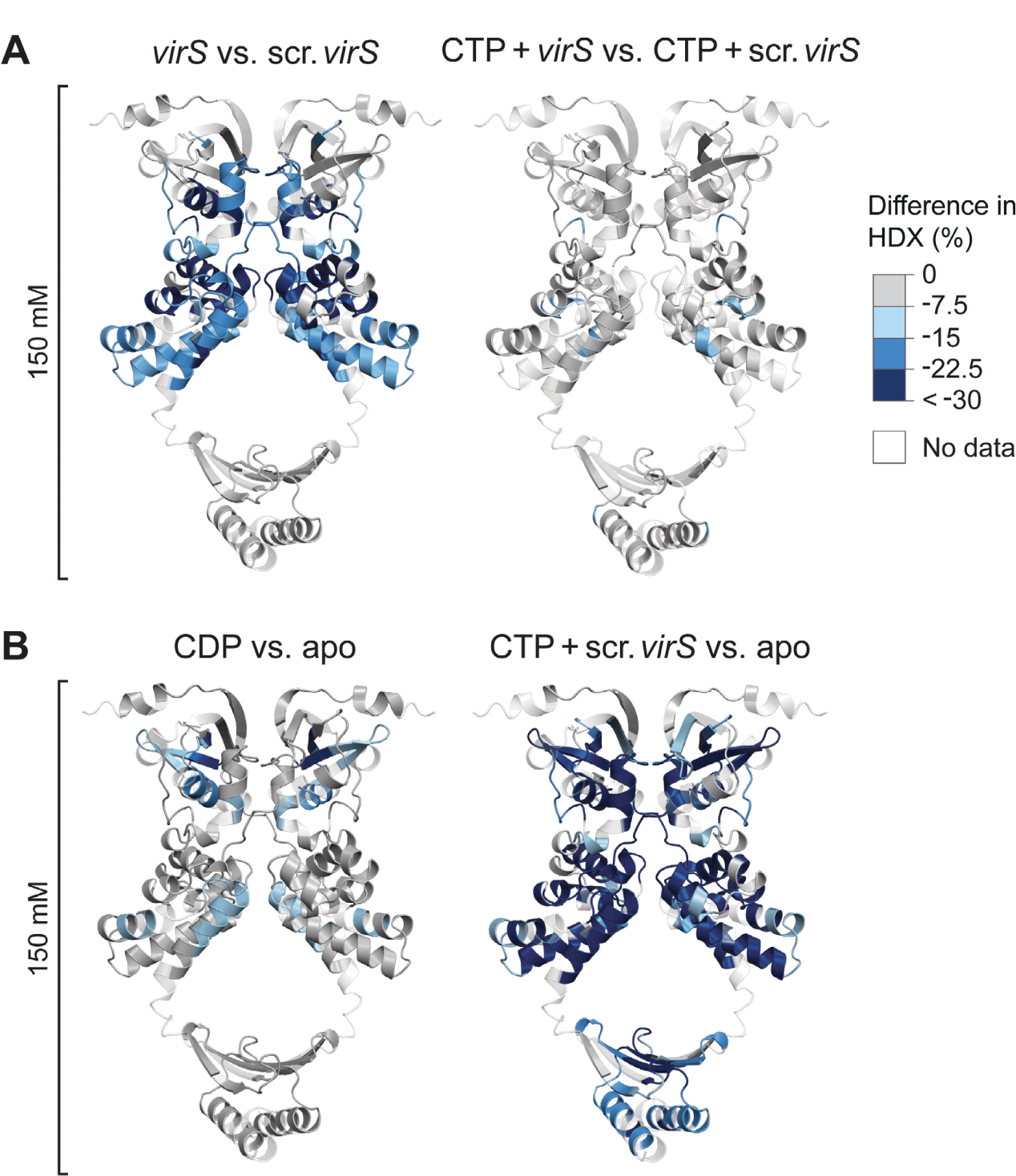
Changes in the HDX pattern of VirB plotted onto the structural model of the VirB dimer. **(A)** Comparison of the HDX patterns of VirB with *virS*-containing DNA compared to scrambled *virS* DNA in the absence and presence of CTP in low-stringency buffer (150 mM NaCl). The color code is given on the right. Blue color denotes reduced HDX in reactions containing *virS* DNA. **(B)** Changes in the HDX pattern of VirB induced by the incubation with CDP or with both CTP and scrambled *virS* DNA compared to the apo state in low-stringency buffer. The color code is given in panel A. Blue color indicates reduced HDX in the ligand-bound state. In both panels, protein regions not covered by any peptides are displayed in transparent white.

**Figure S10.**
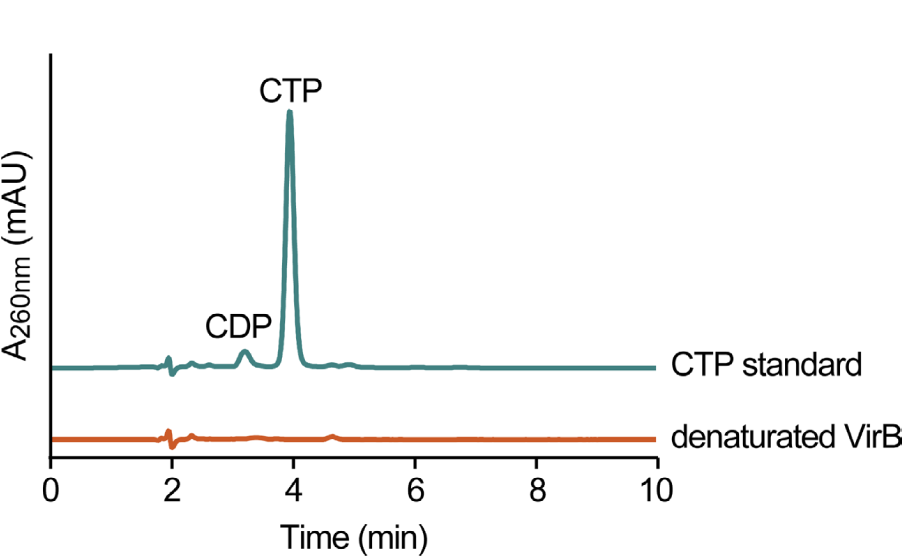
Nucleotide content and CTPase activity of purified VirB. Nucleotide content of VirB. Purified VirB (50 µM) was denatured by the addition of chloroform and heated at 95 °C to release the bound nucleotides. The aqueous phase was then analyzed for the presence of CTP or CDP by HPLC at a wavelength of 260 nm. A CTP standard (100 µM) was analyzed as a reference.

**Figure S11.**
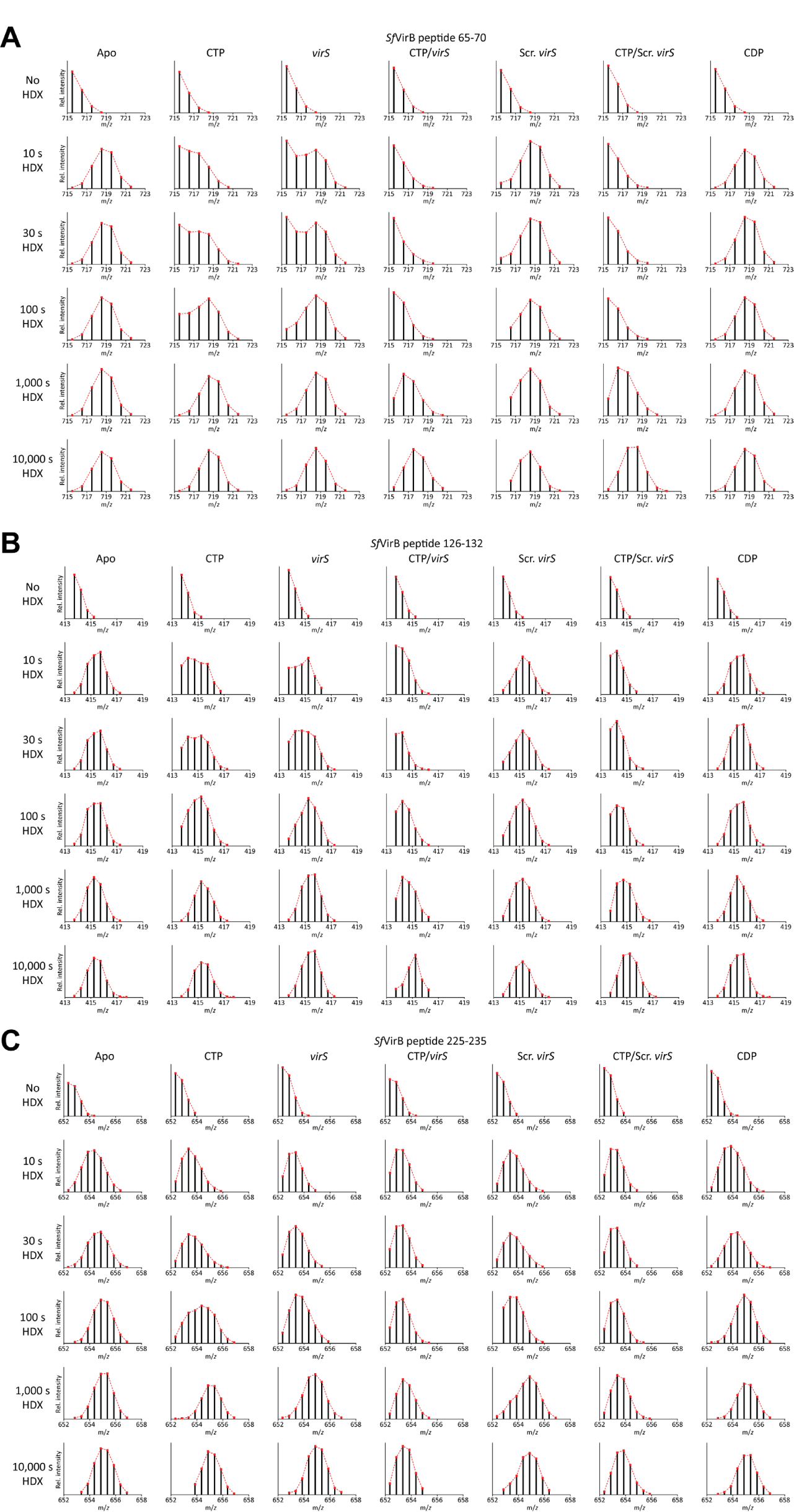
Bimodal HDX behavior of VirB in low-stringency conditions. **(A-C)** Shown are the mass spectra (displayed as peptide ion sticks) of representative VirB peptides obtained after incubation of VirB for the specified time periods in the absence (apo) or presence of the indicated ligands in low-stringency conditions (buffer containing 150 mM NaCl). The distribution of masses indicates EX1 or mixed EX1/EX2 HDX kinetics in some of the conditions. Details of the mass spectra are given in **Data S1**.

**Figure S12.**
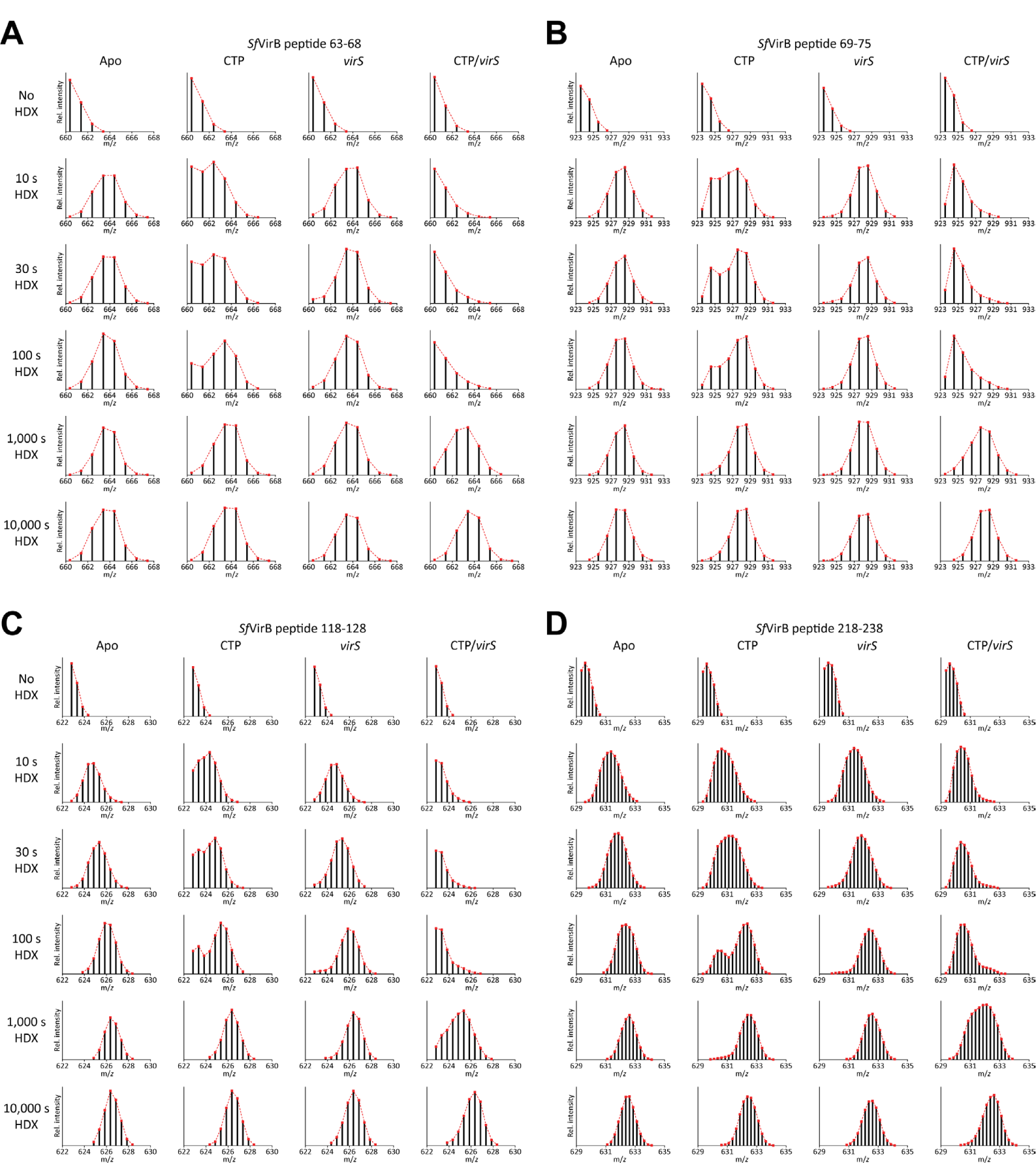
Bimodal HDX behavior of VirB in high-stringency conditions. **(A-C)** Shown are the mass spectra (displayed as peptide ion sticks) of representative VirB peptides obtained after incubation of VirB for the specified time periods in the absence (apo) or presence of the indicated ligands in high-stringency conditions (buffer containing 500 mM NaCl). The distribution of masses indicates EX1 or mixed EX1/EX2 HDX kinetics in some of the conditions. Details of the mass spectra are given in **Data S1**.

**Table S1.**
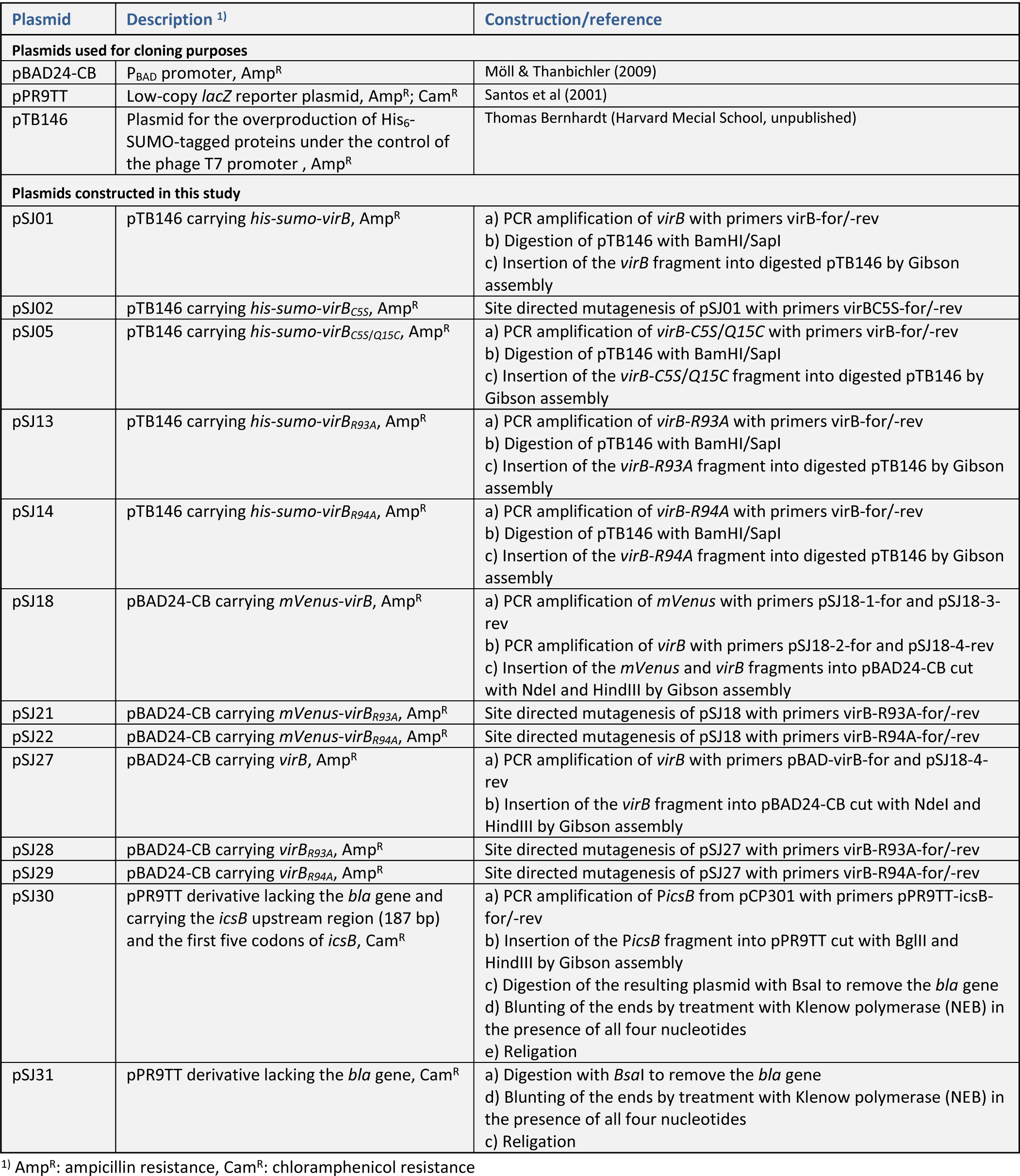
Plasmids used in this study.

**Table S2.**
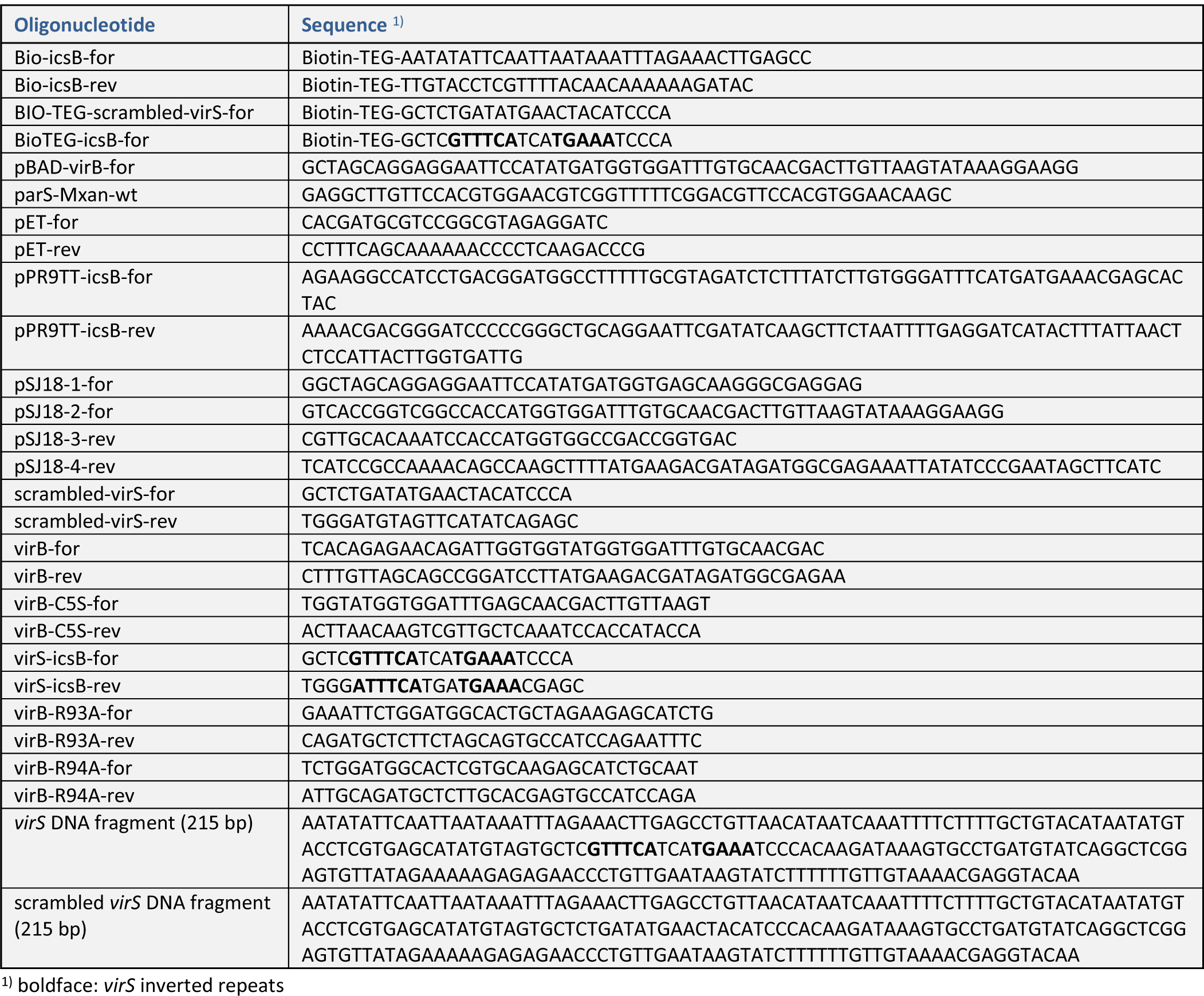
Oligonucleotides and synthetic DNA fragments used in this work.

## Notes

### Competing Interest Statement

The authors have declared no competing interest.

